# StIA is a Novel Nucleoid-Associated Protein of *Streptococcus pneumoniae* that Activates DNA Topoisomerase I

**DOI:** 10.64898/2026.09.11.750838

**Authors:** Antonio A. de Vasconcelos Junior, Pablo Herrera, Adela G. de la Campa, Mónica Amblar

## Abstract

Topoisomerases are essential enzymes that maintain DNA topology. The activity of some of these enzymes is regulated by protein cofactors. Here, we identify a previously uncharacterized nucleoid-associated protein from the human pathogen Streptococcus pneumoniae that binds double-stranded DNA (dsDNA) in a sequence-independent manner both *in vitro* and *in vivo* and activates topoisomerase I (Topo I). This protein was named StIA (Streptococcal Topoisomerase I Activator). *In vivo* analyses showed that deletion of *stIA* impairs bacterial growth in the presence of the Topo I inhibitor seconeolitsine, whereas its overproduction decreases susceptibility to this compound. Consistent with these observations, StIA specifically stimulates Topo I activity without affecting DNA gyrase. StIA forms oligomers and assembles into higher-order nucleoprotein complexes upon DNA-binding. Mechanistic analyses revealed that StIA does not alter DNA cleavage by Topo I but specifically enhances the DNA religation step of its catalytic cycle. This activation is likely mediated through a direct protein-protein interaction, as StIA and Topo I physically associate *in vitro*. Structural modeling further supports the formation of a StIA-Topo I complex and identifies putative contact residues involved in the interaction. Collectively, these findings establish StIA as a novel nucleoid-associated with a dual function: promoting nucleoprotein complex assembly and stimulating Topo I activity through direct interaction with the enzyme. More broadly, our results suggest that the regulation of topoisomerase activity by accessory proteins may represent an important mechanism for fine-tuning DNA topological homeostasis and could provide new opportunities for the development of antibacterial therapies.

## 1. Introduction

To fit inside the cells, the bacterial chromosome must be compacted into a well-organized and dynamic structure known as the nucleoid. This compaction is a fundamental feature of all living organisms, enabling essential DNA transactions such as replication and transcription. In bacteria, chromosome compaction results from the concerted activities of DNA topoisomerases, which maintain DNA at an optimal level of supercoiling (Sc) (Junier et al., 2023), and nucleoid-associated proteins (NAPs) (Dillon and Dorman, 2010), a group of low-molecular-weight proteins that bind to DNA with little or no sequence specificity.

Topoisomerases are essential enzymes that resolve topological constraints by introducing transient DNA breaks. Type I topoisomerases, such as topoisomerase I (Topo I) generate single-strand breaks, whereas type II topoisomerases, including DNA gyrase (gyrase) and topoisomerase IV (Topo IV), introduce transient double-strand-breaks. These enzymes are primary targets of clinically important antibacterial agents such as fluoroquinolones (FQs), which trap type II topoisomerases in covalent cleavage complexes with DNA, thereby blocking replication (Fukuda et al., 2026). Sc regulates transcription, and transcription is, at the same time, a major contributor of the level of Sc. According to the twin Sc-domain model, domains of negative and positive Sc are transiently generated behind and ahead of the transcribing RNA polymerase, respectively (Liu and Wang, 1987). These topological constraints can hinder transcription elongation, and topoisomerases alleviate the resulting torsional stress, thereby ensuring efficient transcriptional progression (Drolet et al., 1994; Massé et al., 1997; Phoenix et al., 1997).

NAPs are necessary for organizing and maintaining the function of bacterial chromosomes. Most possess the ability to alter DNA topology by bending, wrapping or bridging it, thereby modifying its three-dimensional structure (Dorman, 2013). In addition, many NAPs regulate transcription, either positively or negatively (Dillon and Dorman, 2010; Shen and Landick, 2019; Hołówka and Zakrzewska-Czerwińska, 2020), and some have recently been shown to affect topoisomerase activity, highlighting the interplay between chromosome architecture and DNA topology (Ghosh et al., 2014; de Vasconcelos Junior et al., 2023). To date, about twelve NAPs have been identified in Gram-negative bacteria, compared to only six identified in Gram-positives. They differ greatly in abundance, effects and extent of conservation, and show a variety of DNA-binding modes, which illustrate the diversity of their mechanisms of action and roles in chromosome organization. Moreover, extended protein occupancy domains has recently been described in *Escherichia coli* and *Bacillus subtilis* (Freddolino et al., 2021; Amemiya et al., 2022). These multikilobase genomic regions are transcriptionally silent, densely occupied by proteins, and enriched in NAPs, which contribute to gene silencing by excluding RNA polymerase from the DNA.

*Streptococcus pneumoniae* (the pneumococcus) is a devastating human pathogen responsible for a wide range of severe diseases, including community-acquired pneumonia, meningitis, bacteraemia and otitis media. It is estimated to cause approximately one million childhood deaths worldwide each year (WHO, 2007). The widespread emergence of resistance to β-lactams and macrolides has increased the clinical relevance of FQs for the treatment of pneumococcal infections, particularly community-acquired pneumonia (Jacobs et al., 2003; Mandell et al., 2007). Gyrase, Topo IV, and Topo I constitute the complete repertoire of topoisomerases encoded by the pneumococcal genome and are responsible for maintaining chromosome topology (Lucas et al., 2001). Gyrase and Topo I actively regulate Sc, thereby resolving the topological problems associated with DNA replication and transcription (Ferrandiz et al., 2010; de la Campa et al., 2017; García-López et al., 2023b), whereas Topo IV primarily functions in the decatenation of the sister chromosomes after the replication is completed. The role of pneumococcal topoisomerases in Sc homeostasis and its relationship with transcription has been extensively investigated. Transcriptomic analyses of cells subjected to either DNA relaxation, induced by inhibition of gyrase with novobiocin (NOV), or hyper-negative Sc, generated by inhibition of Topo I with seconeolitsine (SCN), revealed a global transcriptional response and uncovered the existence of chromosomal domains with intrinsic topological behavior (Ferrandiz et al., 2010; Ferrándiz et al., 2016; Martín-Galiano et al., 2017). These domains comprise genes with coordinated expression patterns and related biological functions, and can be classified as up-regulated, down-regulated, non-regulated, and AT-rich. In *S. pneumoniae*, Sc homeostasis is regulated primarily through the transcriptional control of topoisomerase genes. DNA relaxation triggers up-regulation of gyrase genes (*gyrA* and *gyrB*) and down-regulation of Topo I (*topA*) and Topo IV (*parC* and *parE*) genes, whereas hyper-negative Sc down-regulates the expression of *topA*. Notably, *topA* and *gyrB* are located within down-regulated (Ferrandiz et al., 2010) and up-regulated Sc domains (Ferrándiz et al., 2014), respectively. Therefore, *topA* transcription closely correlates with changes in Sc density (Ferrándiz et al., 2016), highlighting the central role of Topo I in the Sc homeostasis. Furthermore, pneumococcal Topo I has an essential role in the interplay between transcription and Sc by promoting the formation and stability of the RNA polymerase–DNA complex at promoters during transcription elongation. This function is supported by its physical interaction with RNA polymerase *in vitro* and by their genome-wide proximity *in vivo* (Ferrándiz et al., 2021).

In contrast to the extensive knowledge of pneumococcal topoisomerases, the contribution of NAPs to chromosome organization in *S. pneumoniae* remains poorly understood, and the repertoire of identified pneumococcal NAPs is remarkably limited. Until recently, only HU (Ferrándiz et al., 2018) and SMC (Minnen et al., 2011) have been characterized. HU is the most abundant NAP in this bacterium, is essential for growth and is involved in the maintenance of chromosome architecture. Alterations in HU levels affect Sc *in vivo*, and moderate increments partially counteracts NOV-induced DNA relaxation. By contrast, SMC is involved in chromosome segregation and its deletion results in a mild chromosome segregation defect. More recently, we identified StaR, a novel pneumococcal NAP that regulates Topo I activity (de Vasconcelos Junior et al., 2023). StaR levels are modulated as part of the homeostatic response to DNA relaxation and must remain within a narrow physiological range to sustain normal growth under Sc stress. Consistently, StaR overproduction increases DNA relaxation caused by NOV treatment. Furthermore, StaR co-localizes with the nucleoid, interacts with Topo I *in vivo*, and stimulates its activity. However, despite this recent discovery, the number of characterized NAPs in *S. pneumoniae* remains considerably lower than in other bacterial species, and their contribution to overall chromosome structure is still largely unexplored.

A pneumococcal nucleoid isolation study conducted in our laboratory identified the hypothetical protein Spr0488 as a component exclusively associated with the nucleoid fraction (unpublished results). Spr0488 is a 131 amino acid, highly conserved protein that contains a predicted DNA-binding motif. In this study, we investigated the biological function of Spr0488. *In vivo*, deletion of *spr0488* impaired growth in the presence of the Topo I inhibitor SCN, whereas its overproduction increased resistance to this compound. Biochemical analyses showed that Spr0488 forms dimers, binds dsDNA both *in vivo* and *in vitro* and promotes the formation of high-molecular-weight nucleoprotein complexes upon DNA binding. Furthermore, Spr0488 physically interacts with Topo I and stimulates its DNA relaxation activity *in vitro* by enhancing the latest steps of the catalytic cycle. Collectively, these findings indicate that Spr0488 contributes to the maintenance of Sc and support its classification as a novel nucleoid-associated protein. Accordingly, we propose the name StIA (Streptococcal topoisomerase I Activator) for this newly identified factor.

## 2. Materials and methods

### 2.1. Bacterial strains, plasmids, growth conditions and transformation

*S. pneumoniae* strains used in this study are derivatives of the wild-type strain R6. They were grown at 37 °C as static cultures in a casein hydrolase-based medium (AGCH) supplemented with 0.2% of yeast extract (AY). Carbon sources were normally 0.3% sucrose, and either 0.8% sucrose or 0.8% maltose for strains harbouring pLS1ROM or pStIA plasmids. R6 cells were transformed as described previously (Lacks et al., 1986) and transformants were plated on AY+0.3% or 0.8% sucrose media plates containing 1% agar, which were incubated at 37°C in a 5% CO_2_ atmosphere. *E. coli* was grown in LB medium at 37 °C with shaking. *E. coli* strains DH5α and BL21-CodonPlus (DE3) (Agilent Technologies, Santa Clara, CA, USA) were transformed according to Hannahan (Hanahan, 1983). Bacterial growth was followed by measuring OD_620nm_ either with a visible spectrophotometer (1100 Spectrophotometer, Fisher Bioblock Scientific) or a microplate reader (Infinite F200, Tecan).

The pneumococcal *stIA* deletion mutant (Δ*stIA*) was constructed by insertion-deletion of a kanamycin resistance cassette (Km^r^) through allelic replacement mutagenesis (Song et al., 2005). For this purpose, three fragments were generated by the polymerase-chain reaction (PCR) using Phusion Hot Start HiFi (Thermo Fisher). Two of them, which contain the upstream and downstream *stIA* regions, were obtained using oligonucleotide pairs 0488up_F/0488Up_R_Kpn and 0488Down_F_Bam/0488Down_R, respectively (Table 1). The third fragment, containing Km^r^, was amplified from plasmid pR410 (Sung et al., 2001) using primers Km_Kpn and Km_Bam. The three resulting PCR products were purified, digested with BamHI and KpnI, and ligated. The ligation mixture was further amplified using 0488up_F and 0488Down_R primers and the resulting fragment used to transform strain R6 (Figure 1A). Transformants were selected with 250 µg/ml of Km. Recombinant clones were confirmed by PCR amplification with external oligonucleotides 0488_F_ext and 0488_R_ext, and Sanger sequencing.

**Figure 1.**
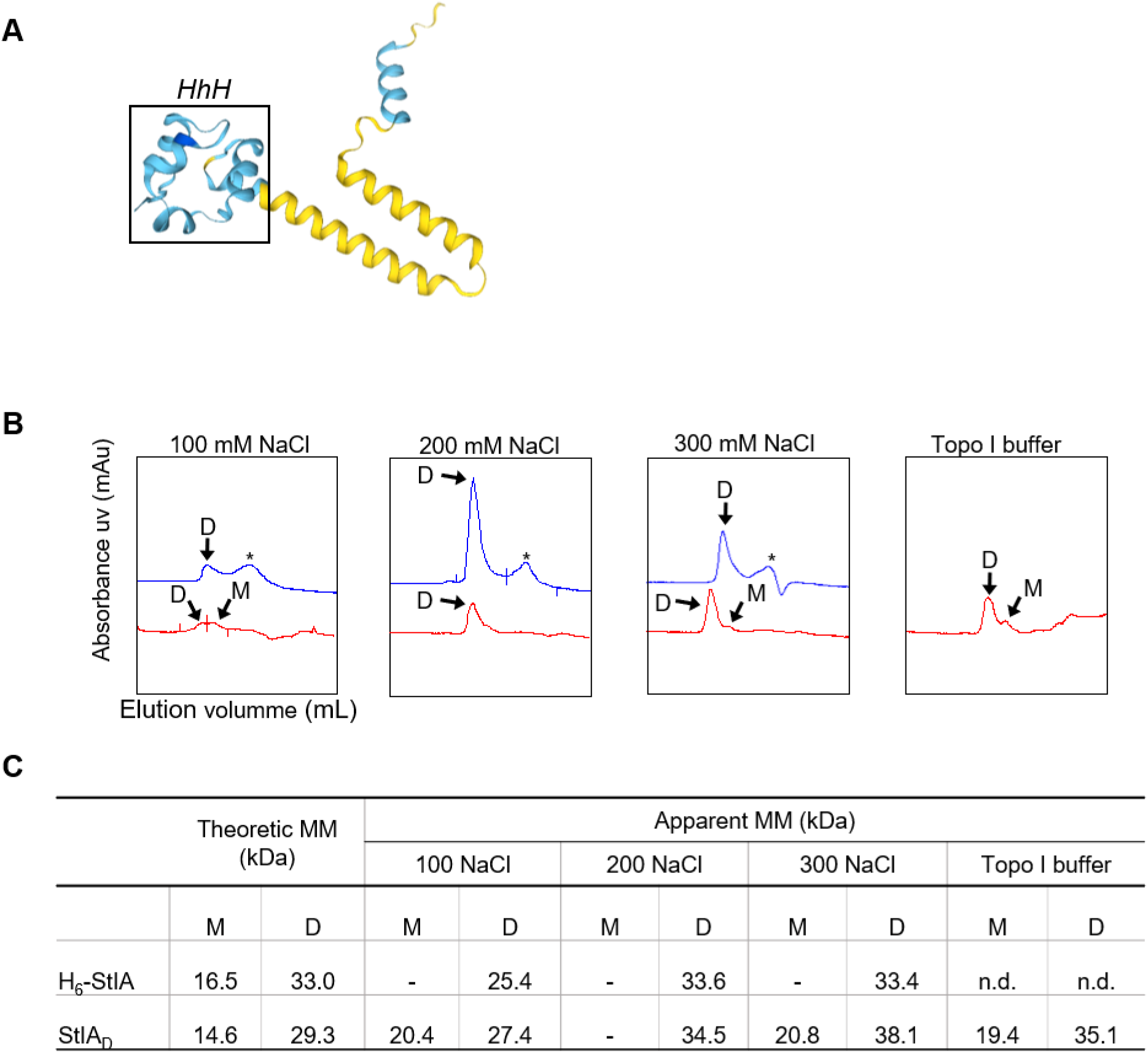
StIA possess a DNA-binding domain and is mainly a dimer. **(A)** Tertiary structure of StIA was predicted by SWISS-MODEL. Residues are coloured by their local quality value using QMEANDisCo (range 0-1). Confidence class code: dark blue (score>0.9) very high confidence; light blue (0.7 >score> 0.9) confident; yellow (0.5 >score> 0.7). The HhH domain is framed within a box. **(B)** Multimerization analysis of H_6_-StIA and StIA_D_ performed by size exclusion chromatography as described in the Materials and methods section in different saline buffers. Chromatograms of H_6_-StIA (blue) and StIA_D_ (red) with diverse eluents containing the indicated NaCl concentrations, or the buffer used in Topo I *in vitro* DNA relaxation assays, are shown. Arrows indicate peaks corresponding to the dimer (D) or the monomer (M). Other peaks corresponding to lower molecular mass particles are labelled by an asterisk. Elution volume (ml) of each peak was used to estimate the apparent molecular mass (MM) by plotting in the column calibration curve. **(C)** Table with theoretic and apparent MM of the monomer (M) and dimer (D) of H_6_-StIA and StIA_D_.

**Table 1.**
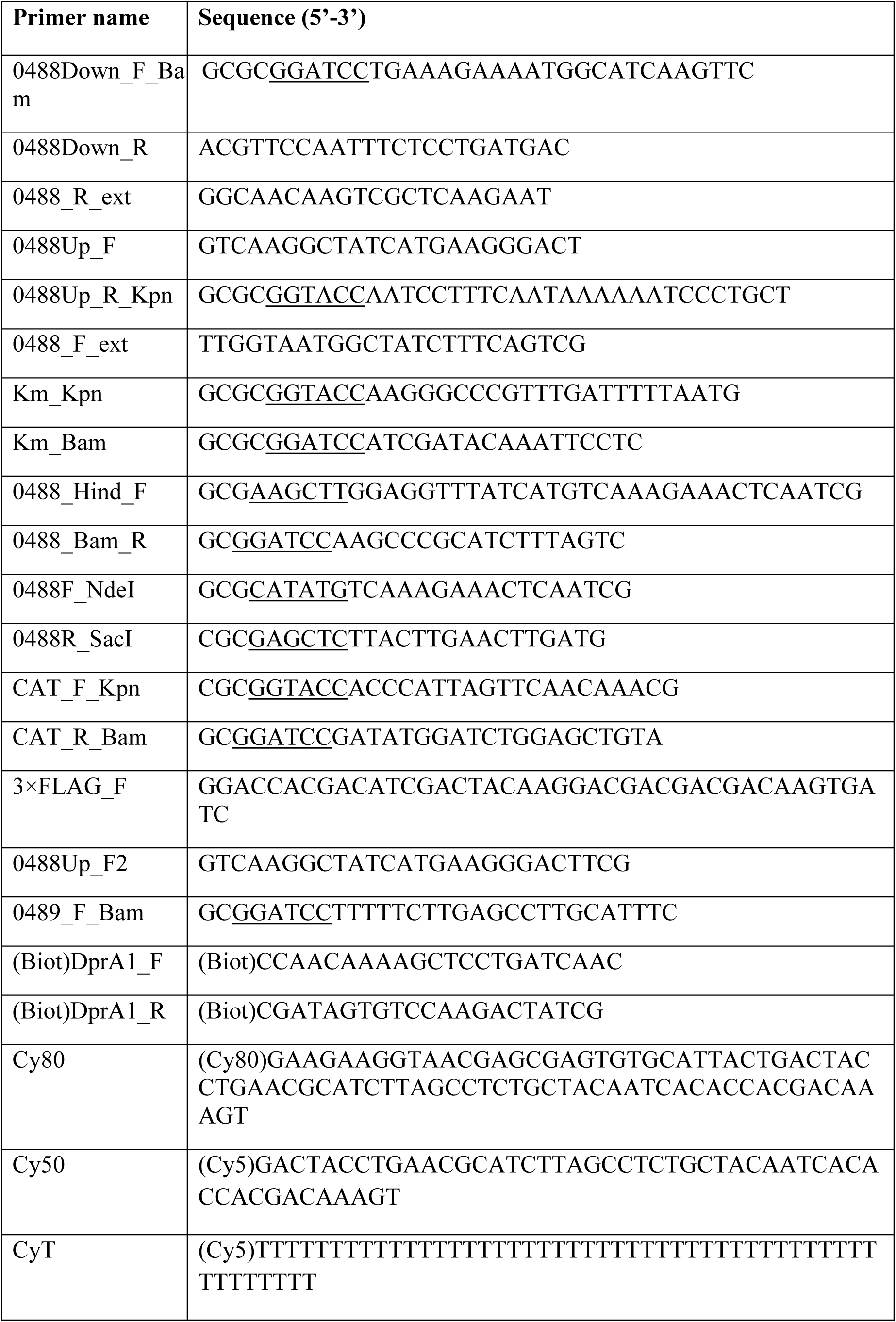
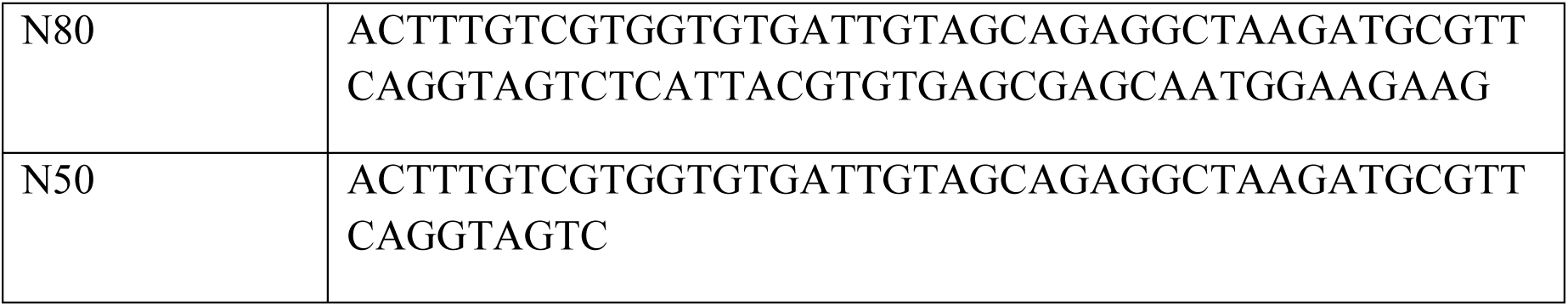
List of primers used in this study. The underlined sequences indicate enzyme-restriction sites.

The pStIA plasmid was obtained by cloning a 599-bp fragment containing *stIA* (including its ribosome-binding site) into pLS1ROM (Ruiz-Masó et al., 2012) under the control of the P_Mal_ promoter. The *stIA*-containing fragment was amplified by PCR from R6 chromosomal DNA using primers 0488_Hind_F and 0488_Bam_R (Table 1), digested with BamHI and HindIII and cloned into vector pLS1ROM cut with the same enzymes. The resulting recombinant plasmid pRStIA was then introduced into strain Δs*tIA* by transformation and selected with 1 µg/ml of erythromycin to obtain strain Δ*stIA* [pStIA]. Cloning was verified by sequencing. As control strains, R6 and Δ*stIA* were transformed with pLS1ROM, thus obtaining R6 [pLS1ROM] and Δ*stIA* [pLS1ROM] strains. Plasmid pET28a-StIA encoding a 6×His-tag fused to the N-terminus of StIA was obtained by cloning *stIA* into the expression vector pET28a. A PCR fragment containing *stIA* was amplified from R6 genomic DNA with primers 0488F_NdeI and 0488R_SacI, digested with NdeI and SacI, and cloned into pET28a cut with the same enzymes. Plasmid pET28a-StIA was transferred to *E. coli* BL21-CodonPlus (DE3) (Agilent Technologies).

### 2.2. Overexpression of StIA and protein purification

*E. coli* BL21-CodonPlus (DE3) [pET28a-StIA] was grown in LB medium supplemented with 50 µg/ml kanamycin at 37 °C to OD_600nm_ = 0.4. Expression of H_6_-StIA was induced by addition of 1 mM isopropyl thio-β-D-galactoside (IPTG) at 30 °C and samples were withdrawn at various times. To analyse the overproduction of StIA, samples were suspended in SLB 4× (Bio-Rad), supplemented with 0.5 M β-mercaptoethanol, lysed by incubation at 100 °C for 5 min and loaded in 4-20% SDS-polyacrylamide gel. Solubility of target protein was analysed as previously described (Amblar et al., 1998). Briefly, cell pellets were suspended in 50 mM Tris–HCl, pH 7.5, 2 mM EDTA, 0.1% Triton X 100 and 0.1 mg/ml lysozyme, and lysed by incubation at 37 °C for 30 min. The viscosity of the extracts was reduced by passage through a 0.36 mm inner diameter needle. Extracts were centrifuged, and samples of the supernatant (soluble form) and pellet (insoluble form) were examined by 4-20% SDS-polyacrylamide gel electrophoresis. To purify H_6_-StIA, 2 l of the strain grown as described above were induced with 1 mM IPTG at 30 °C for 1h. Cells were harvested by centrifugation, washed with phosphate-buffered saline (PBS) and suspended in 64 ml of buffer A (50 mM Tris-HCl pH 8.0, 0.5 M NaCl). Cells were disrupted by sonication and extracts clarified by centrifugation as previously described (de Vasconcelos Junior et al., 2023). Imidazole was added up to 40 mM, and the crude extract was applied to a 5 ml HiTrap column (Cytiva) equilibrated with buffer A containing 40 mM imidazole. Protein elution was achieved in an ÄKTA prime system (GE Healthcare) with 110 ml of a 40 to 400 mM imidazole gradient in buffer A, and fractions were analysed by 4-20% SDS-polyacrylamide gel electrophoresis. Imidazole was removed from samples by dialysis against 20 mM Tris-HCl pH 7.8, 1 mM EDTA, 0.2 M NaCl, and glycerol was added to the sample to get 50% before storing at -20 °C. Protein concentration was determined with a Qubit 4 fluorimeter using the Qubit® Protein Assay Kit (Invitrogen).

Topo I, GyrA and GyrB were purified by affinity chromatography in a Ni-NTA (Qiagen) column as described previously (Balsalobre et al., 2011).

### 2.3. Thrombin cleavage of His-Tag

The His_6_-tag fused to StIA was removed by digestion with Thrombin (Novagen, supplier Merck, Millipore) using 0.5 units of biotinylated thrombin per mg of purified H_6_-Spr488 at room temperature for 16 h, following instructions from manufacturers. Biotinylated Thrombin was then removed from reaction using the Thrombin Cleavage Capture kit. Digestion was confirmed by electrophoresis in a Criterion^TM^ TGX 4-20% polyacrylamide gels (Bio-Rad) and Coomassie staining.

### 2.4. Size exclusion chromatography

The oligomerization state of H_6_-StIA prior and after digestion with thrombin (StIA_D_) was analyzed by size exclusion chromatography with a Superdex 200 10/30 GL column (Cytiva) using an ÄKTA prime plus. A calibration curve was performed using the Gel Filtration Calibration kit LMW (Cytiva) plus alcohol dehydrogenase (Sigma), using Tris buffers (50 mM Tris pH8) with different ionic strength: either 100, 200, or 300 mM NaCl, and the Topo I activity buffer (50 mM Tris pH8, 10 mM MgCl_2_ and 1 mM DTT and 100 mM KCl). H_6_-StIA and StIA_D_ were dialyzed against the column buffer to remove glycerol. A total sample volume of 100 µl containing 5.3 mg/ml of each protein was loaded at a flow rate of 0.5 ml/min. Protein elution was followed by measurement absorbance at 280 nm. Molecular weights were estimated by plotting the peak elution volume on the calibration curve.

### 2.5. Western blotting

Whole cell lysates (∼5 × 10^5^ cells) were obtained by centrifugation of 10 ml cultures (OD_620nm_ = 0.4). They were suspended in 400 μl of SLB 4× (Bio-Rad, Hercules, CA, USA) supplemented with 0.5 M β-mercaptoethanol and incubated for 5 min at 100 °C. Lysates were separated on Any kD™ Criterion™ TGX Stain-Free™ Protein Gels (Bio-Rad) and transferred to 0.2 μm PVDF membranes with a Trans-Blot Turbo Transfer System (Bio-Rad) at 25 V and 1 A for 30 min. Membranes were blocked with 5% skim milk in Tris-buffered saline overnight and incubated with anti-Topo I (1:500), anti-GyrA (1:2000), anti-RpoB (1:500) and anti-StIA (diluted 1:2000) for 1 h. Rabbit polyclonal antibody against StIA was obtained from Davids Biotechnologies with 2 mg of purified H_6_-StIA extracted from a SDS-polyacrylamide gel following a 63-day immunization protocol. Anti-rabbit IgG-peroxidase Ab (Sigma-Aldrich-Merck) was used as the secondary antibody. SuperSignalWest Pico chemiluminescent substrate (Thermo-Fisher) was used to develop the membranes. Signal was detected with a ChemiDoc^TM^ MP system (Bio-Rad) and images were analyzed using Image Lab^TM^ software (Bio-Rad). Molecular masses of GyrA and TopoI are 92 kDa and 79 kDa, respectively.

### 2.6. *In vitro* cross-linking assays

For multimerization analysis of StIA, cross-linking assays were performed as previously described (Ferrándiz et al., 2018) with some modifications. Briefly, proteins were incubated in the presence of 0.1% glutaraldehyde at room temperature for 30 min in 20 µl of 10 mM Tris pH 8, 150 mM NaCl, 25 % glycerol, 0.25 mM EDTA, 0.25 mM DTT. Samples were diluted with 6 µl of Laemmli Sample Buffer 4 ’ and loaded in a 4-20% SDS-polyacrylamide gel. Gels were stained with Coomassie blue for 1 h. To study the physical interaction between StIA and Topo I, 100 pmol of StIA (1.5 µg) and 50 pmol of Topo I (4 µg) were incubated at room temperature for 5 min in 40 µl of PBS 1× with 1 mM DTT and 0.002% of glutaraldehyde. Reactions were terminated by addition of 250 mM Tris-HCl, pH 8.0 and solubilized with an equal volume of Laemmli Sample Buffer 2×. Electrophoresis was conducted in 4-20% SDS-polyacrylamide gels and cross-linked proteins were analysed by western blotting using anti-StIA (diluted 1:2000) as described above.

### 2.7. Nucleoid isolation

*S. pneumoniae* R6 cells were grown in 40 ml of AY *+* S to OD_620nm_ = 0.4 and were harvested by centrifugation. The pellet was resuspended in 1 ml of PBS 1 × supplemented with 10 µl of Protector RNase inhibitor (Roche). Then 1 ml of PBS 1× with 1% deoxycholate was added and incubated at room temperature for 1 h. The suspension was filtered with 0.45 µm filter and 500 µl were loaded on a Superose® 6 increase 10/300 GL column (Cytiva) equilibrated in 25 mM Tris pH 6.8, 50 mM NaCl, 5% glycerol and connected to and ÄKTA Pure 25 chromatography system. The column was previously calibrated with blue dextran (2000 kDa), thyroglobulin (669 kDa), ferritin (440 kDa), aldolase (158 kDa), conalbumin (75 kDa), carbonic anhydrase (29 kDa) and ribonuclease A (13.7 kDa) as molecular weight standards. Fractions of 1 ml were collected. To assess the DNA and protein elution profile, samples were analysed by measuring absorbance at 260 nm in a microplate reader (Infinite F200, Tecan) and the Qubit® Protein Assay (Molecular Probes], respectively.

### 2.8. DNA binding assays

Binding of H_6_-StIA and StIA_D_ to dsDNA was analysed through electrophoretic mobility-shift experiments (EMSAs) on a 261 bp fragment (5’ biotin-labelled) obtained by PCR using R6 chromosome as template, the oligonucleotide pair (Biot)DprA1_F/ (Biot)DprA1_R as primers (Table 1), 200 µM dNTPs and 1.25 units of Taq Platinum DNA polymerase (Thermo Fisher). The binding assays were performed in a 100 µl volume containing 200 fmol of DNA, 20 mM Tris-HCl pH 7.8, 100 mM NaCl, 2 mM MgCl_2_, and different amounts of either H_6_-StIA or StIA_D_. The mixture was incubated at 37 °C for 30 min and the binding was terminated by addition of 11 μl of 30% glycerol, 1% xylene cyanol, 1% bromophenol blue and 10 mM EDTA. Reactions (30 μl) were fractionated in 5% Criterion TBE (89 mM Tris-base, 89 mM Boric acid, pH 8.0, 2 mM EDTA) polyacrylamide Gels (Bio-Rad) at 150 V for 1.8 h and 4 °C. DNA was transferred to an Hybond-N+ membrane (Amersham) by electroblotting in 1 × TAE buffer (40 mM Tris, 20 mM Acetic acid, pH 8.0, 1 mM EDTA) at 35 V for 2 h, using the Trans-blot cell from Bio-Rad. The blots were UV-crosslinked in a CL-1000 UV Crosslinker (UVP, Inc.) by irradiation at 254 nm and 120 mJ and detection was performed using a Chemiluminescent Nucleic Acid Detection Module (Thermo Fiscer) following manufacturer’s instructions. Bands were visualized with a ChemiDoc MP System (Bio-Rad).

Binding to fluorescent substrates was also performed. Single and total or partially double stranded substrates were generated by combining the fluorescent labelled oligonucleotides (5’ cyanine-5-labelled, Cy): 80-nt oligonucleotide (Cy80), 50-nt oligonucleotide (Cy50); or homopolymer of 50 thymines (CyT); with the full or partially complementary non-labelled oligonucleotides: N80 (of 80-nt in length) or N50 (of 50-nt in length). The O3 DNA substrate with a 3’ overhang of 30-nt was generated by annealing of the Cy50 with the non-labelled N80; the O5 substrate with a 5’ overhang of 30-nt was generated by annealing of the Cy80 with the N50 oligonucleotide. To generate the O3 and O5 substrates, 50 nM of each oligonucleotide, were mixed in 10 mM Tris-HCl pH8, 50 mM NaCl and incubated at 95 °C for 2 min. The mixture was slowly cooled to 25 °C in a thermocycler with a raping rate of 1.3% during 40 min and incubated for 5 additional min at the same temperature, followed by a rapid cooled down to 4 °C. Binding assays were performed by incubating 10 nM of each substrate in 20 µl volume containing 20 mM Tris-HCl pH 8, 150 mM NaCl, 2 mM MgCl_2_, 1 mM DTT, and serial dilutions of protein, for 20 min at 37°C. To have a total protein concentration of 0.0075 mg/ml in all reaction tubes, the precise amount BSA was added to each mixture. Four µl of Blue Juice sample loading buffer (Invitrogen) were added to the reactions and they were separated in 5% Criterion TBE polyacrylamide gels at 100 V and 4 °C for 90 min. Gels were visualized in a ChemiDoc MP.

Binding assays to pBR322 was carried out in 15 µl volume containing 1 nM of the indicated plasmid form, 20 mM Tris pH8, 100 mM KCl, 1mM DTT, 2 mM MgCl_2_, 0,014 mg/ml BSA and different amounts of StIA_D_. Mixtures were incubated at 30°C for 20 min and binding was terminated by addition of 6 μl of 6 × Loading Dye (Thermofisher Scientific). Samples were run in a 1× TBE 0.5% agarose gel at 18 V for 14-15 h. Gels were stained with 0.5 µg/ml of ethidium bromide and visualized with UV light. Linearized plasmid (L) with 3’-cohesive-ends was generated upon PaeI digestion. Nicked plasmid (nc) was generated by digestion with the nickase Nt.BstNBI, which introduces 4 nicks on pBR322. Negatively supercoiled (CCC) and relaxed pBR322 were obtained from Inspiralis.

### 2.9. DNA topoisomerase I and gyrase activity assays

Assays of pBR322 relaxation by Topo I were carried out as described previously (de Vasconcelos Junior et al., 2023). Briefly, 1.8 nM (500 ng) of CCC pBR322 were incubated with purified Topo I at 37 °C during 1 h in 200 µl of activity buffer (20 mM Tris-HCl pH 8, 100 mM KCl, 10 mM MgCl_2_, 1 mM DTT, 50 µg BSA/ml). When indicated, either H_6_-StIA or StIA_D_ were added. Reactions were terminated by 2 min incubation with 50 mM EDTA and treated with 1% SDS, 100 µg/ml proteinase K for 1 h at 37 °C. Samples were analysed by electrophoresis in 1 ×TBE 1% agarose gels run at 18 V for 18 h. After electrophoresis, the gel was stained with ethidium bromide and bands corresponding to the CCC, OC and RC plasmid forms were quantified using the Image Lab program (Bio-Rad laboratories). Topo I activity was calculated as the percentage of both, the appearance of topoisomers plus the OC + RC forms, and the disappearance of the CCC form.

Plasmid nicking assays were performed incubating 400 ng of pBR322 and different amounts of StIA_D_ in 100 µl of cleavage buffer (activity buffer without MgCl_2_ and with 1 mM EDTA). When indicated, DNA was pre-incubated with Topo I for 15 min at 4 °C to allow binding, before addition of StIA_D_. Reactions were incubated at 37 °C for 1 h, treated as described above, and resolved in 1 × TBE 1.2% agarose gel. Gels were stained with ethidium bromide and the intensity of nicked and CCC plasmids was determined using image Lab software.

Cleavage and religation assays were performed using the 5’-Biotin-labelled 32-mer STS-oligonucleotide previously described. DNA cleavage was carried out by incubating Topo I in the cleavage buffer in the presence of increasing concentrations of StIA_D_, for 1 h at 37 °C to yield a 5’-end-labelled 19-mer cleavage product. The reactions were terminated with 45 % formamide dye and heating at 95 °C for 5 min. The samples were resolved on 15% denaturing PAGE using a 1× TBE as a running buffer at 200 V and transferred to Hybond-N+ membrane (Amersham) in 1×TAE buffer at 30 V for 16 h. The blots were UV-crosslinked and bands were detected and visualized as described earlier. To determine the effect of StIA_D_ in Topo I mediated intramolecular religation, a cleavage reaction was performed for 1 h followed by the incubation with StIA_D_ for 5 min. The religation reaction was initiated by addition of 10 mM MgCl_2_, incubated for additional 1h and terminated by 10 min incubation at 37 °C with 50 µg/ml proteinase K. The reaction products were resolved in 15% denaturing PAGE.

Enzymatic reactions with gyrase were conducted as described previously (Fernández-Moreira et al., 2000).

### 2.10. Structural modelling and docking

A dimeric structural model of StIA (M0) was obtained using the Multimer mode of the ClusPro protein–protein docking server. ClusPro performs rigid-body sampling followed by clustering of low-energy solutions and supports homo-multimer modelling through its Multimer functionality (Kozakov et al., 2017; Ashizawa et al., 2026). The resulting M0 model was used as the common structural scaffold for subsequent interaction analyses.

For protein–protein docking, the M0 StIA model was independently docked using pyDock and ClusPro against pneumococcal DNA topoisomerase I (Uniprot database annotation number Q8DPI9). pyDock performs rigid-body docking using FTDock and ranks candidate orientations using primarily electrostatic and desolvation contributions together with a van der Waals term (Jiménez-García et al., 2013). ClusPro uses rigid-body sampling followed by clustering of low-energy conformations and ranks models based on cluster representation and energy (Kozakov et al., 2017; Ashizawa et al., 2026; Comeau et al., 2004). The three highest-ranked solutions from each method were retained for comparative structural analysis. Docked structures were visualised analysed using PyMOL open-source version and UCSF ChimeraX 1.12. Intermolecular residue contacts were defined as atom–atom distances ≤4.0 Å. Atom pairs separated by <2.0 Å were classified as potential steric clashes and used as a geometric quality-control criterion. Solvent-accessible surface areas were calculated with a solvent radius of 1.4 Å and dot density of 2 with solvent exposure enabled, following the solvent-accessible surface approach. Buried surface area was calculated according to Belapure et al (Belapure et al., 2023). Residue-level consensus was assessed by determining the frequency with which individual StIA residues occurred in the interfaces of independent docking models. For the principal StIA– Topo I consensus, the three pyDock and the first two ClusPro models were considered. The third ClusPro solution showed an alternative model involving both StIA monomers and a different geometry.

## 3. Results

### 3.1. StIA is a highly conserved protein that displays a typical DNA binding motif

A homology search using *S. pneumoniae* R6 StIA as a query revealed that all pneumococcal genomes encode highly conserved orthologs sharing more than 96% sequence identity. BLASTP analysis further identified homologous proteins in other *Streptococci* with sequence identity higher than 87%. Analysis of the UniProt database classified StIA as a member of the Rad51 N-terminal domain-like superfamily (SSF47794), characterized by the presence of a helix-hairpin-helix (HhH_5) domain. This domain is located at the N-terminus of the eukaryotic DNA repair protein Rad51 (Aihara et al., 1999) and at the C-terminus of the *E. coli* transcriptional regulator NusA (Eisenmann et al., 2005).

Structural motif prediction using SMART (Letunic and Bork, 2026) and InterPro (Blum et al., 2025), confidently identified some distinct features: a disordered region between residues 1 and 20, a low-complexity plus a coiled-coil region spanning residues 24 and 78, and an HhH domain located between residues 78 and 126 (E-value of 3.88 × 10^-2^). Consistent with these predictions, a three-dimensional structure model generated using the automated SWISS-MODEL server (https://swissmodel.expasy.org/) predicted a similar overall architecture, with a Global Model Quality Estimate (GMQE) of 0.73 (Figure1A). Among the predicted structural elements, the HhH_5 domain displayed the highest confidence score. This compact domain is a conserved structural module found in a wide range of prokaryotic and eukaryotic DNA-binding proteins, including nucleases, polymerases, helicases, DNA repair enzymes, and transcriptional terminators. Consists of a bundle of 4 to 5 α-helices arranged as two orthogonally packed α-hairpins and contains one classic and one pseudo HhH motif. Each motif comprises two α-helices connected by a short loop and mediates non-sequence-specific DNA binding through interactions with the DNA phosphate backbone (Doherty, 1996).

The predicted coiled-coil region consists of two or more α-helices wrapped around each other to form a supercoiled structure. Coiled-coil motifs primarily mediate protein– protein interactions and are frequently found in proteins that assemble into dimers, trimers, or higher-order oligomers, thereby contributing to oligomerization and overall structural stability (Cabezon et al., 2000; Szczepaniak et al., 2021).

### 3.2. Free StIA is a dimeric protein in solution

Multimerization is a common feature of small DNA-binding proteins, and structural predictions suggested that StIA adopts a coiled-coil structure. To determine its oligomerization state, StIA was overexpressed in *E. coli* BL21(DE3) using the pET28b expression system and purified as an N-terminal His_6_ fusion protein (H_6_-StIA). The His_6_ tag was subsequently removed by thrombin digestion, generating StIA_D_, which retains only three additional residues. The oligomeric state of both the H_6_-StIA and StIA_D_ was analysed by size exclusion chromatography under different salt concentrations. Elution profiles were compared with molecular weight standards to estimate their apparent molecular masses (Figure 1B-C). At 200 and 300 mM NaCl, both proteins eluted as a single narrow peak, indicating highly homogeneous samples. The apparent molecular mass of H_6_-StIA (33.6 and 33.4 kDa at 200 and 300 mM NaCl, respectively) closely matched the theoretical mass of a globular dimer (33.0 kDa), and no monomeric species were detected. StIA_D_ also eluted predominantly as a dimer, although with slightly higher apparent molecular masses (34.5 and 38.1 kDa). At 300 mM NaCl, a minor peak of approximately 20.8 kDa, consistent with a monomer, was also observed.

At 100 mM NaCl, both proteins displayed broader, bimodal elution profiles, indicating increased molecular heterogeneity. Under these conditions, the apparent molecular masses of the major species decreased to 25.4 for H_6_-StIA and 27.4 kDa for StIA_D_. This behaviour likely reflects either a reduced hydrodynamic radius or non-specific electrostatic interactions with the chromatographic matrix, consistent with the basic nature of the protein. In the Topo I activity buffer, containing 100 mM KCl and 10 mM MgCl_2_, StIA_D_ exhibited an elution profile comparable to that observed at higher NaCl concentrations, with a major peak of 35.1 kDa and a minor monomeric peak of 19.4 kDa. Such observation may be due to subtle changes in the conformational flexibility of charged surfaces induced by the presence of Mg^2+^ and /or the different electrostatic and hydrodynamic properties of the K⁺ ion.

The apparent molecular masses of StIA_D_ were consistently higher than both the theoretical values and those of H_6_-StIA, suggesting that removal of the His_6_ tag slightly alters hydrodynamic properties. These values are consistent with predominantly dimeric protein adopting a non-globular conformation and containing a minor monomeric fraction. Collectively, these results support the conclusion that free StIA exists predominantly as a dimer in solutions at salt concentrations ≥100 mM. Furthermore, the N-terminal His_6_ tag does not affect dimer formation but appears to promote a more compact protein conformation.

### 3.3. Levels of StIA affect susceptibility to the Topo I inhibitor SCN

The previous identification of StIA in nucleoids isolated from *S. pneumoniae* prompted us to investigate its contribution to chromosome structure and Sc homeostasis. To this end, a deletion mutant (Δ*stIA*) was generated by allelic replacement with a Km^r^ cassette, as described under material and methods (Figure 2A). In parallel, *stIA* was cloned into the pLS1ROM vector under the control of the maltose-inducible promoter P_Mal_, yielding plasmid pStIA, which was introduced into the *ΔstIA* strain to generate the complemented strain *ΔstIA* [<u>p</u>S<u>tIA</u>] (Figure 2B).

**Figure 2.**
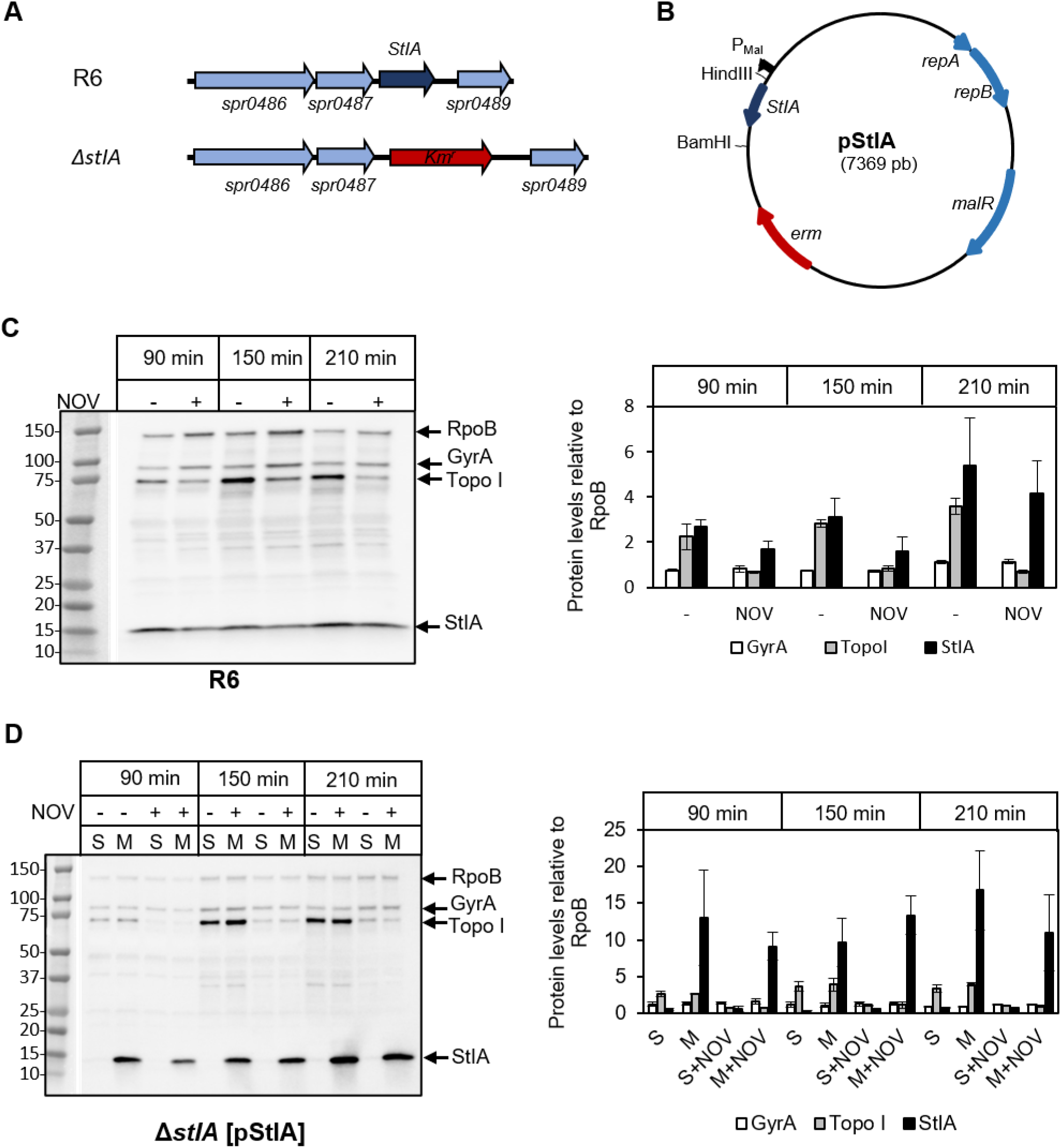
The levels of StIA are not affected by DNA relaxation induced by NOV treatment. **(A)** *stIA* genomic context of R6 and *ΔstIA* strains, in which deletion of *stIA* was carried out by insertion of Km^r^. Genes are shown as arrows. **(B)** Structure of the recombinant pStIA plasmid that carries *stIA*. Genes involved in replication (*repA* and *repB*), the one coding for the P_Mal_ repressor (*malR*), and the erythromycin resistance (*erm*) gene are indicated. (C, D) Effect of NOV on StIA levels. Cultures of R6 **(C)** or *ΔstIA* [pRStIA] (D) were grown to OD_620 nm_ = 0.3, diluted 200-fold in medium with 0.3% sucrose for R6 or with 0.8% of sucrose (S) or maltose (M) for *ΔstIA* [pStIA] and either with (+) or without (-) 0.5 × MIC of NOV. Samples taken at 90, 150 and 210 min were analysed by Western blotting (left panels) and levels of the indicated proteins were quantified (right panels). Antibodies against GyrA (1:2000), Topo I (1:500), StIA (1:2000) and RpoB as internal control (1:2000) were used. Levels of GyrA, Topo I and StIA relative to RpoB are shown. Data are the average of three independent replicates ± SEM.

StIA expression was analysed by Western blotting in both the wild type strain (R6) and *ΔstIA* [pStIA] strain. StlA levels were compared with those of GyrA and Topo I, using RpoB as internal control (Figure 2C and D). Cultures were grown in the presence or in the absence of NOV at 0.5 × MIC, and samples were taken after 90, 150 and 210 min of treatment. In strain R6, StIA levels remained essentially unchanged throughout the experiment regardless of NOV treatment, indicating that its expression is not responsive to changes in Sc (Figure 2C). This observation is consistent with previous transcriptomic analyses showing that *stIA* expression is unaffected by gyrase inhibition (Ferrandiz et al., 2010). In strain *ΔstIA* [pStIA], the levels of StIA were dependent on induction with maltose, increasing between 3- and 6-fold relative to R6 strain at all times tested. As previously reported (de Vasconcelos Junior et al., 2023), NOV treatment reduced the levels of Topo I in both R6 (between 3.5- and 4.2-fold) and *ΔstIA* [pStIA] (between 3.3- and 5.2-fold), whereas GyrA levels remained unchanged. Importantly, neither Topo I nor GyrA abundance was influenced by StIA overproduction, indicating that StIA does not regulate topoisomerase expression.

To determine whether StIA contributes to Sc homeostasis, the growth of R6, Δ*stIA*, Δ*stIA* [pStIA], and the control strain Δ*stIA* [pROM], which carries the pLS1ROM vector, was evaluated in the presence of drugs altering Sc, either via gyrase (NOV) or Topo I (SCN) inhibition. As expected, treatment of the R6 wild type strain with either antibiotic at 0.5 × MIC impaired growth. Deletion of *stIA* did not affect growth under standard conditions or in the presence of NOV compared to R6, as reflected by growth curves and duplication times (Fig. 3A). In contrast, clear differences were observed during growth in the presence of SCN. Although the doubling times of R6 (76.7 ± 3.9 min) and Δ*stIA* strains (83.1 ± 6.0 min) were not significantly different, the Δ*stIA* strain reached a lower maximum optical density and entered the lysis phase earlier than R6. Quantification of the area under the curve confirmed a significant reduction in growth of the Δ*stIA* strain during SCN treatment (312 ± 6.5 for R6 versus 236 ± 8.2 for Δ*stIA*).

**Figure 3.**
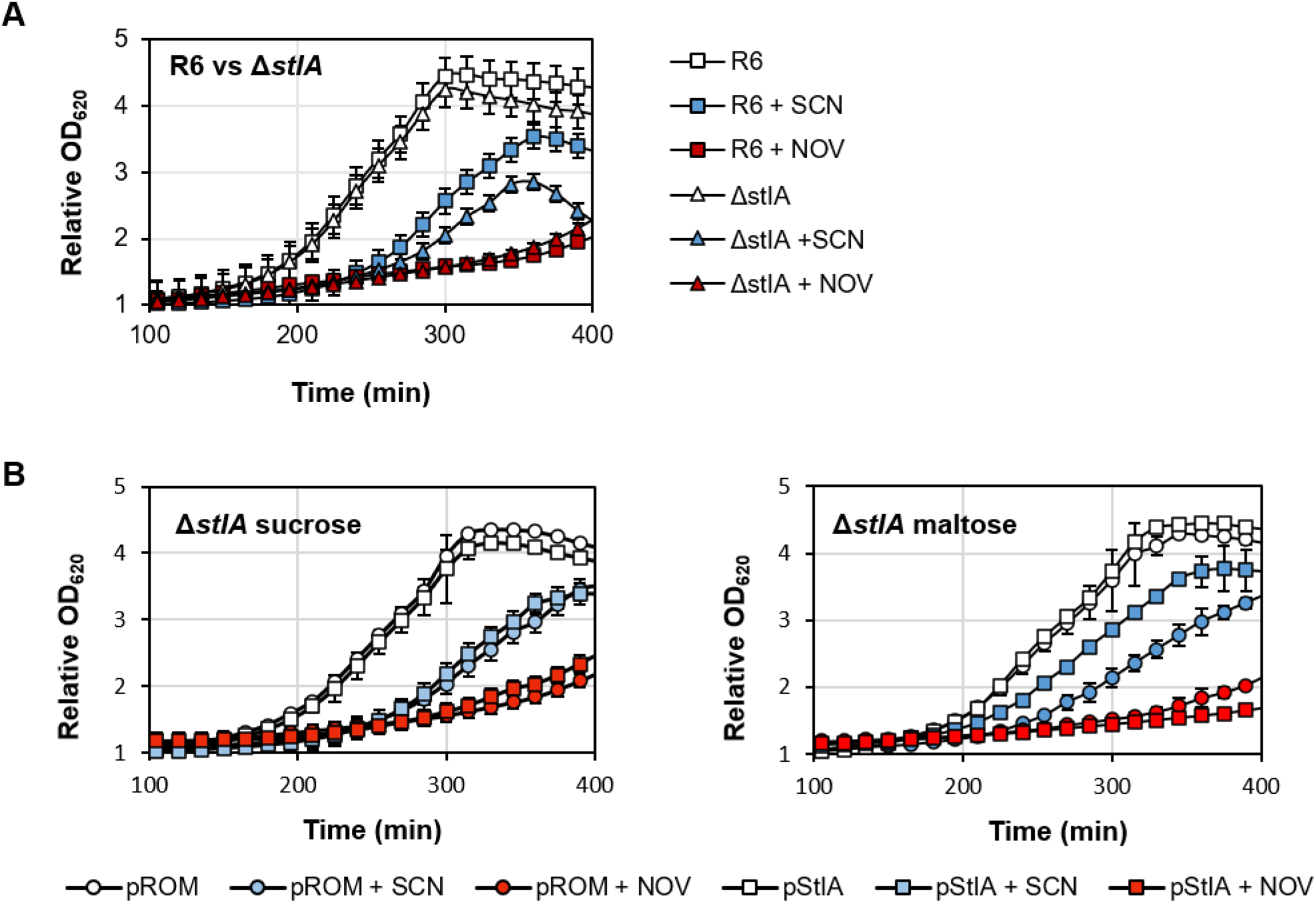
Effect of SCN or NOV treatment on bacterial growth upon *stIA* deletion or overexpression. Cultures were grown without antibiotic to OD_620_ = 0.3, they were then diluted 200-fold in media with 0.5 × SCN (blue) or 0.5 × NOV (red), or without antibiotic (white). **(A)** Growth of strains R6 and Δ*stIA* in 0.3 % sucrose. **(B)** Growth of strains Δ*stIA* [pROM] and Δ*stIA* [pStIA] in the presence of either 0.8 % sucrose or 0.8% maltose. Growth was monitored in a TECAN Infinite 200 PRO reader. Data are the average of three independent replicates ± SEM.

The effect of increased StIA levels was also examined using the inducible Δ*stIA* [pStIA] strain. In the presence of sucrose, which represses *stIA* expression, Δ*stIA* [pStIA] and Δ*stIA* [pROM] strains displayed indistinguishable growth under all conditions tested (Fig. 3B). Induction of *stIA* with maltose had no detectable effect on growth in the absence or presence of NOV. In contrast, StIA overproduction significantly improved growth in the presence of SCN (Figure 3B). Under these conditions, Δ*stIA* [pStIA] exhibited a shorter doubling time (mean ± SD of 87.0 ± 2.4 min) than the control Δ*stIA* [pROM] strain (108.4 ± 10.9 min), indicating that a 3- to 5-fold increase in StIA levels partially alleviates the growth defect caused by Topo I inhibition. Collectively, these results demonstrate that cellular StIA levels specifically influence susceptibility to SCN, but not to NOV, supporting a functional role for StIA in Sc homeostasis through modulation of Topo I activity.

### 3.4. StIA stimulates Topo I activity but has no effect on gyrase

To determine whether StIA modulates Topo I, we examined the effects of both H_6_-StIA and StIA_D_ on the *in vitro* Topo I activity (Figure 4). Relaxation of plasmid pBR322 by Topo I was measured in the presence of increasing concentrations of either H_6_-StIA or StIA_D_. Enzymatic activity was quantified by measuring both, the increase in relaxed topoisomers and the corresponding decrease in the negatively Sc covalently closed circular (CCC) form. Because the electrophoretic conditions used did not allow discrimination between open circular (OC) and relaxed circular (RC) forms, these species were quantified together with topoisomers of intermediate linking number as a measure of Topo I activity.

**Figure 4.**
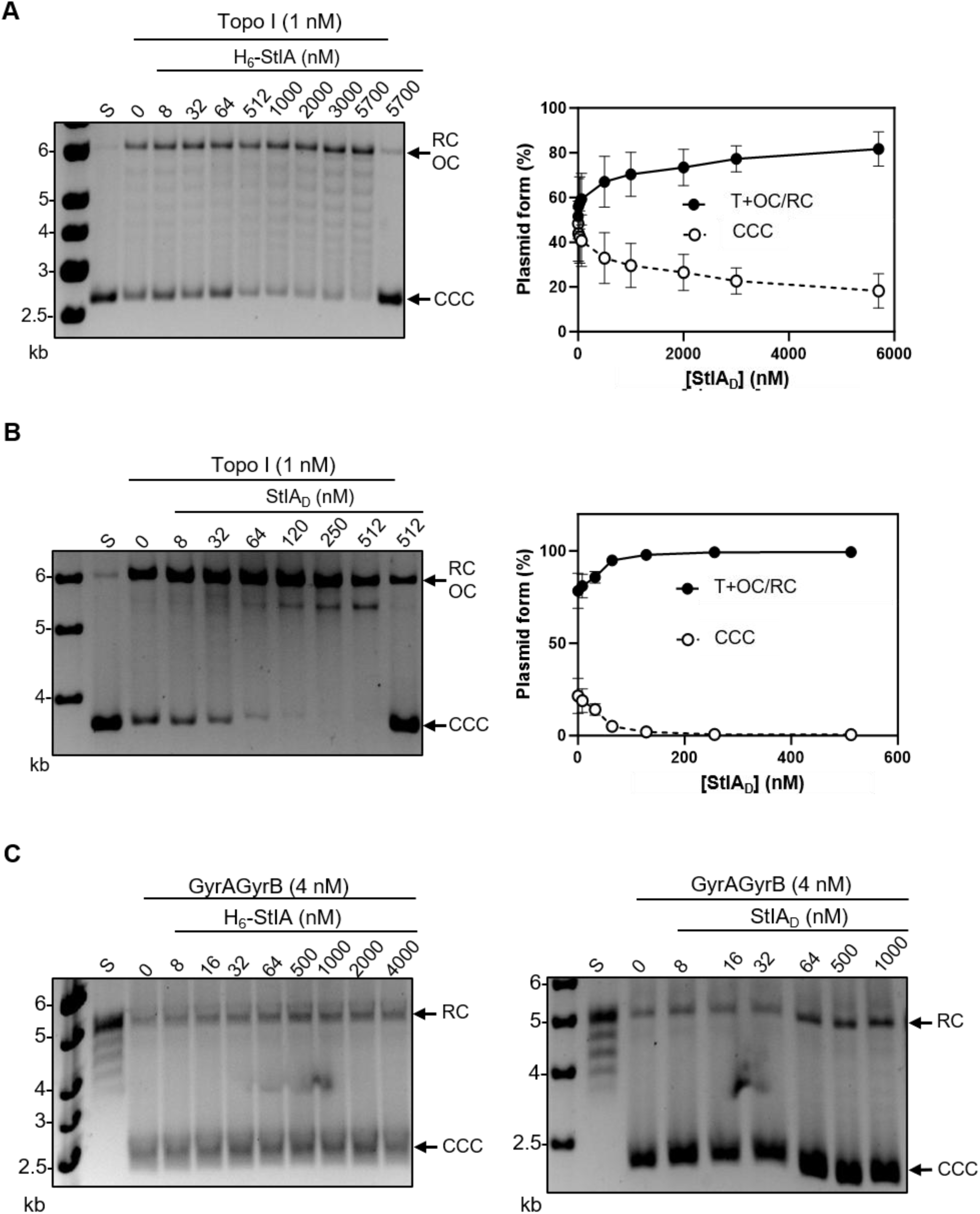
StIA specifically activates Topo I relaxation activity. Relaxation of pBR322 by Topo I in the presence of H_6_-StIA **(A)** or StIA_D_ **(B)**. Plasmid was incubated at 37 °C with Topo I for 1 h, and StIA was added at the indicated concentrations. Samples were processed as described in the Materials and methods section and analysed in 1 % agarose gels (left panel). Left wells, molecular weight standards in kb; S, substrate; CCC, negatively supercoiled covalently closed form; OC, open circles; RC, relaxed circular plasmids. Topo I activity determined as the increase of the topoisomers (including OC/RC forms) and decrease of CCC in the presence of H_6_-StIA or StIA_D_ is shown (right panel). Results are the average ± SEM of three independent replicates. **(C)** Effect of H_6_-StIA (left panel) or StIA_D_ (right panel) on gyrase activity. Supercoiling activity assays with 4 nM of reconstituted gyrase were performed in the presence of the indicated StIA concentrations as described in methods and analysed in 1 % agarose gel.

The results showed that both proteins stimulated Topo I-mediated relaxation of pBR322. In the presence of H_6_-StIA, the ratio of topoisomers plus OC/RC DNA increased from 51.6 ± 3.2 % in the absence of protein to 81.7 ± 3.6 % at the highest concentration tested, with a concomitant decrease in the CCC from 48.4 ± 3.2 % to 18.3 ± 3.6 % (Figure 4A). Likewise, StIA_D_ increased the ratio of topoisomers plus OC/RC from 78.5 ± 9.5% in the absence of protein to 99 % ± 0.3 % at the highest concentration, accompanied by the corresponding reduction in the CCC form (Figure 4B). Neither H_6_-StIA nor StIA_D_ exhibited intrinsic relaxation activity when tested alone at concentrations equal to or higher than those used in Topo I assays (the rightmost wells on Figure 4A and 4B), demonstrating that the observed increase in DNA relaxation resulted from stimulation of Topo I rather than from an independent catalytic activity of either protein. Notably, stimulation by StIA_D_ was detected at concentrations as low as 32 nM, whereas 512 nM of H_6_-StIA was required to produce a comparable effect, indicating that the digested protein is a substantially more potent activator of Topo I. These results suggest that the fusion of 20 residues at the N-terminus significantly impairs the ability of StIA to stimulate Topo I activity.

The ability of StIA to modulate DNA gyrase activity was also evaluated. DNA supercoiling assays were performed using reconstituted gyrase and relaxed pBR322 as substrate in the presence of either H_6_-StIA or StIA_D_ (Figure 4C). Neither the proportion of CCC DNA nor that of the OC/RC species changed with increasing concentrations of either protein, indicating that StIA does not affect DNA gyrase activity under the conditions tested.

### 3.5. StIA binds to dsDNA in vitro and in vivo

The stimulation of Topo I activity by StIA could result from StIA-induced changes in the DNA substrate that enhance its accessibility to Topo I. Because structural analyses predicted that StIA contains a canonical DNA binding motif, its DNA-binding activity was investigated by electrophoretic mobility shift assays (EMSA).

Initially, DNA binding was assessed using a 261-bp biotin-labelled dsDNA generated by PCR as substrate for either H_6_-StIA or StIA_D_ (Figure 5A). H_6_-StIA showed retarded bands at concentrations of 256 nM (corresponding to a protein-to-DNA binding stoichiometry of 128:1), consistent with the formation of DNA-protein complexes (Fig. 5A). Incubation with StIA_D_ at concentrations of ≥256 nM resulted in complete retention of the DNA in the wells, with no detectable free DNA, suggesting the formation of high-molecular-weight DNA-protein complexes. The abrupt transition from no detectable binding to complete retention further suggests that multiple StIA_D_ molecules can bind cooperatively to dsDNA. These results demonstrated that both proteins bind dsDNA and indicate that StIA_D_ displays a higher apparent DNA-binding capacity than H_6_-StIA.

**Figure 5.**
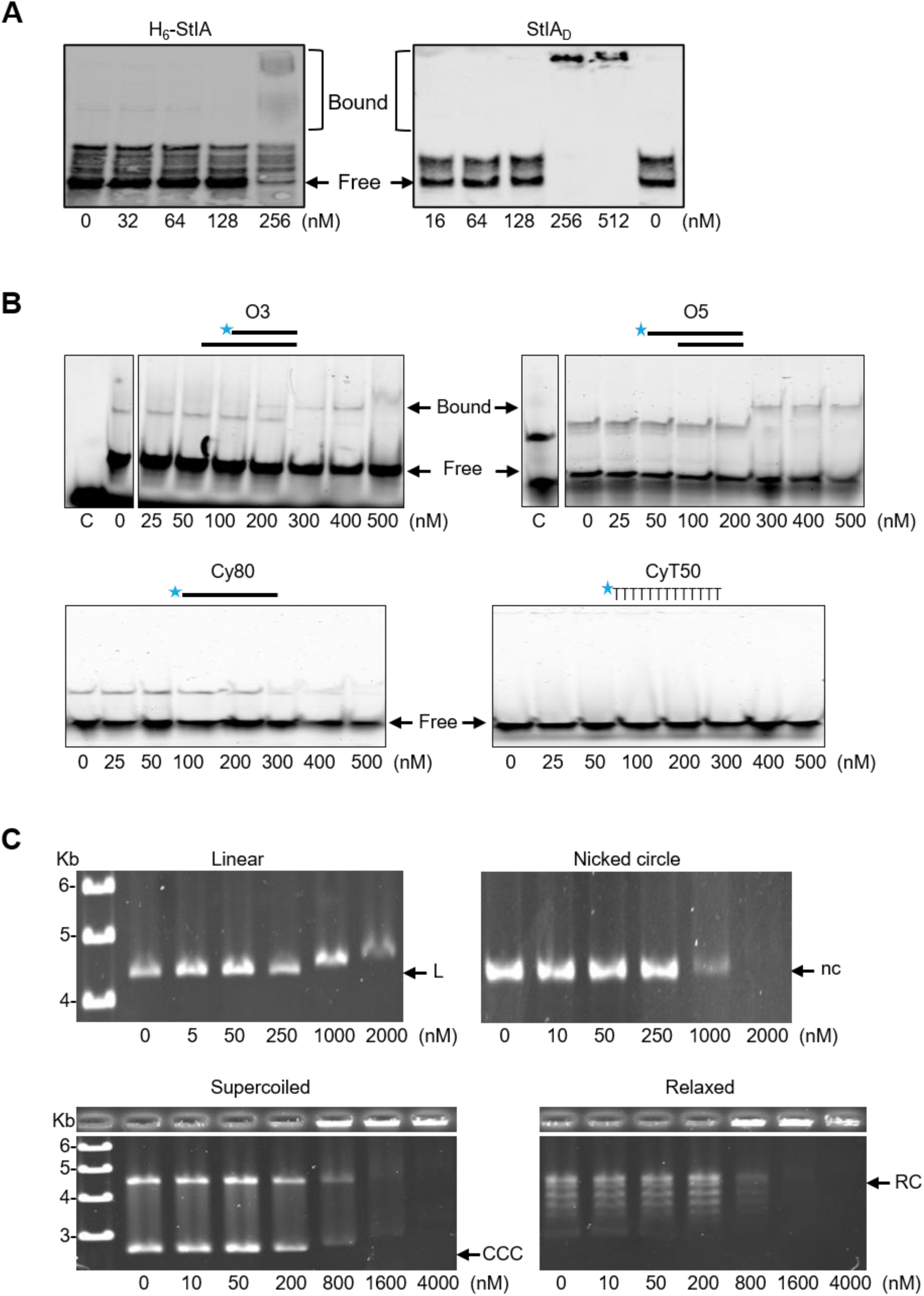
StIA binds dsDNA but not ssDNA. **(A)** Binding of H_6_-StIA and StIA_D_ to a 261-bp PCR biotin-labelled fragment. Reactions were performed by incubating 2 nM of substrate with the indicated concentrations of protein at 37 °C for 30 min. **(B)** Binding of StIA_D_ to fluorescent labelled substrates. 10 nM of the following substrates: O3, O5, Cy80 or CyT50 were incubated with the indicated concentrations of protein for 20 min at 37°C. Reactions were analysed in a 1 × TBE 5%-polyacrylamide non-denaturing gel and visualized in a Chemidoc apparatus. **(C)** Binding of StIA_D_ to pBR322. 1 nM of plasmid linearized with PaeI (L), nicked circle generated by digestion with nuclease Nt.BstNBI (nc), CCC plasmid or relaxed plasmid (RC), were incubated with the indicated concentrations of protein for 20 min at 30°C. Reactions were analysed in a 1 × TBE 0.5% agarose gel and stained with 0.5 µg/ml of ethidium bromide. For CCC and RC plasmids well retention is shown.

To minimize the formation of these large complexes and better characterize substrate specificity, shorter fluorescently labelled oligonucleotides were used. Fluorescent 5’- end labelled 50- and 80- mer oligonucleotides were annealed to fully or partially complementary strands to generate a panel of DNA substrates, including ssDNAs (Cy80 and CyT), fully dsDNA, and partially dsDNAs containing either a 3’ (O3) or a 5’ (O5) single stranded overhang. These substrates were incubated with increasing concentrations of StIA_D_ and analysed by EMSA (Figure 5B). StIA_D_ bound efficiently to fully and partially dsDNA, whereas no binding was detected with either single stranded substrates, regardless of sequence or length. Binding efficiencies for O3 and O5 substrates were comparable, with a protein-to-DNA binding stoichiometry of approximately 30:1. Equivalent binding efficiencies were observed for H_6_-StIA (nor shown), indicating that the N-terminal His_6_ tag does not affect DNA binding affinity or substrate specificity when short oligonucleotide substrates are used.

During DNA relaxation, Topo I binds negatively Sc CCC DNA, transiently generates a nicked intermediate, and subsequently religates the DNA to produce topoisomers with progressively lower linking numbers until the RC form is generated. To determine whether StIA preferentially recognizes specific plasmid topoisomers or reaction intermediates, its binding to linear (L), nicked circular containing 4 nicks (nc), CCC, and RC forms of pBR322 was examined. Each DNA was incubated with increasing concentrations of StIA_D_ and binding was assessed by changes in electrophoretic mobility shift on agarose gels (Fig. 5C). Retarded DNA bands were observed for all plasmid forms at protein concentrations from 800 nM to 1 µM, demonstrating that StIA_D_ binds all plasmid forms with similar efficiency, irrespective of its structure or topology. Again, DNA bands diminished intensity or completely disappeared at the highest protein concentration, primarily with CCC and RC plasmids, which were largely retained in the loading wells. These results are consistent with the formation of high-molecular-weight nucleoprotein complexes likely resulting from the binding of numerous StIA molecules per plasmid (binding stoichiometry of 2000:1). Moreover, they confirmed that StIA_D_ is a double-stranded DNA-binding protein capable of associating extensively throughout the DNA *in vitro* regardless of its sequence, structure, or topology.

Previous isolation of pneumococcal nucleoids indicated that StIA is associated with the nucleoid (unpublished results), suggesting that StIA may also bind DNA *in vivo*. To confirm this, nucleoids were separated from the cytoplasmic fraction of R6 cell extracts by gel filtration chromatography. Cells were gently lysed, chromosomal DNA was partially fragmented, and the lysate was separated by gel filtration. Nucleoid and cytoplasmic fractions were identified according to their elution profile and absorbance at 260 nm, and total DNA and protein content were quantified (Figure 6). The elution pattern of StIA, was analysed by Western blotting and the blots were subsequently re-probed with antibodies against the prototype nucleoid associated protein HU. StIA co-eluted with the DNA-containing high-molecular-weight fractions corresponding to nucleoids, being eminently as an oligomeric protein. A comparable distribution was obtained with HU, in which monomer and high-order oligomers co-fractionated with the DNA, consistent with its association to the nucleoid. These findings suggest that StIA is specifically associated with the pneumococcal nucleoid *in vivo* and, together with the extensive DNA-binding observed *in vitro*, support the conclusion that StIA is a nucleoid-associated protein that likely associates with chromosomal DNA as a multimeric complex.

**Figure 6.**
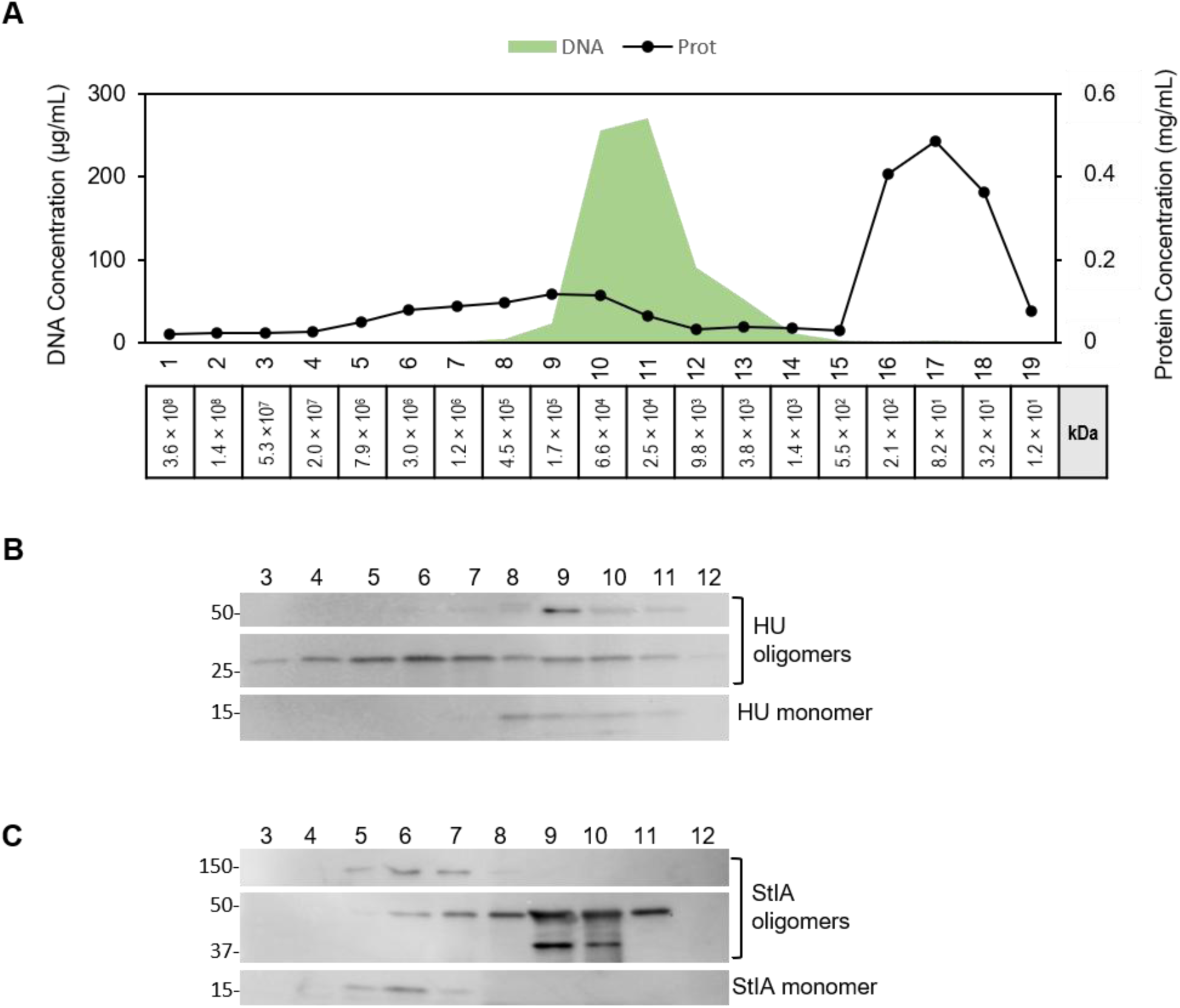
StIA localizes to the nucleoid. **(A)** Fractionation of cell extracts from R6 cultures by gel filtration. Fractions of 1 ml were collected, and their DNA (green) and protein content (black) was measured as described in the Materials and methods section. Apparent molecular masses (in kDa) of particles eluted on each fraction estimated according to their Ve are shown. Western blotting of collected fractions using antibodies against **(B)** StIA (1:500) or **(C)** HU (1:100). MW marker in kDa is shown on the left.

### 3.6. StIA and Topo I physically interact in vitro

The specific stimulation of Topo I activity by StIA suggested the possibility of a direct interaction between the two proteins. To test this hypothesis, chemical cross-linking experiments were performed using the purified proteins. Topo I and StIA_D_ were incubated together in the presence of glutaraldehyde, and the resulting protein complexes were analysed by Western blotting (Figure 7). Immunodetection with the anti-StIA antibody revealed a band with an apparent molecular mass of 100 kDa (Figure 7A). This band was absent when StIA_D_ was incubated alone and is consistent with the expected molecular mass of a StIA_D_ – Topo I complex (93.5 kDa), providing evidence for a direct interaction between the two proteins *in vitro*. In addition, bands corresponding to StIA_D_ dimers, trimers and higher-order oligomers were detected upon glutaraldehyde treatment, further confirming the oligomerization capacity of StIA observed above. Faints bands migrating at approximately the same molecular mass as Topo I were also detected with the anti-StIA antibody, most likely due to cross-reactivity with the His_6_-tag present in the purified Topo I preparation.

**Figure 7.**
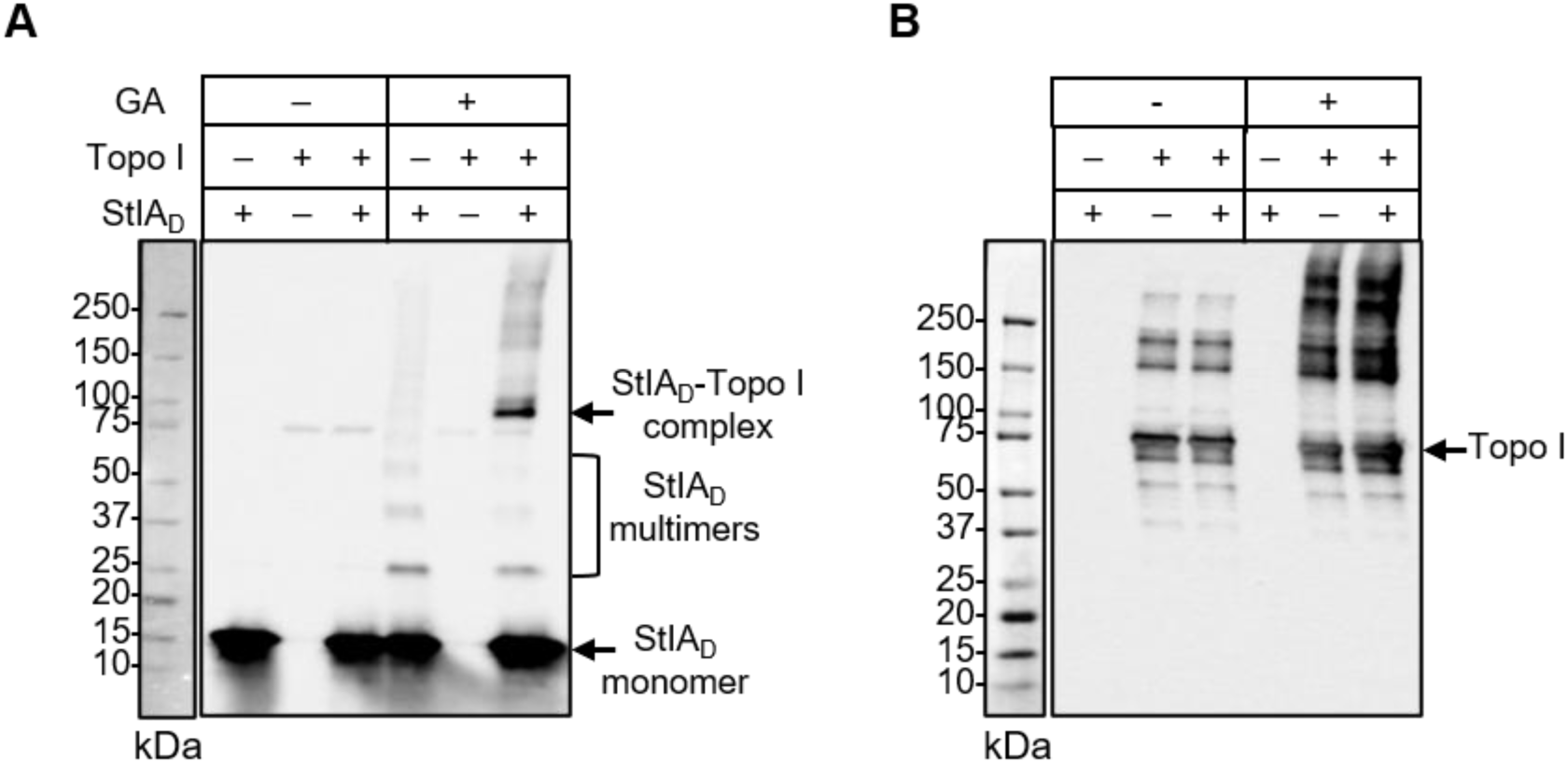
StIA physically interacts with Topo I *in vitro*. *In vitro* crosslinking of purified Topo I and StIA_D_. 50 pmol of Topo I and 100 pmol of StIA_D_ were incubated at room temperature with 0.002% glutaraldehyde (GA) as described in the Materials and methods section. Reactions were loaded in 4-20% SDS-polyacrylamide gels and bands were detected by Western blotting using antibodies anti-StIA **(A)** or the anti-Topo I **(B)**. Bands corresponding to StIA_D_, Topo I and StIA_D_-Topo I complex are indicated. The molecular weight standard is shown on the left.

When the membrane was probed with anti-TopoI antibody, multiple high-molecular-mass cross-linked species were detected, precluding the unequivocal identification of a putative StIA-Topo I complex (Figure 7B).

### 3.7. StIA stimulates intramolecular religation catalysed by Topo I but not DNA cleavage

After binding to DNA, Topo I introduces a single-strand cleavage, passes the intact strand through the nick, and catalyses the subsequent intramolecular religation. Some type IA topoisomerases require Mg^2+^ for religation but not for DNA cleavage (Tse-Dinh, 1986; Bhat et al., 2009), allowing each step to be analysed independently. To elucidate the mechanism of Topo I activation by StIA, we first investigated its Mg^2+^ requirement in both cleavage and religation. Different Topo I amounts were incubated with CCC pBR322 in the absence or in the presence of 10 mM MgCl_2_, and cleavage was measured as the percentage of nicked plasmid. In the absence of MgCl_2_, a unique band corresponding to the nicked plasmid increases with Topo I concentration, indicative of Topo I cleavage activity (Figure 8A). Maximum cleavage (34.4% of nicked plasmid) was achieved after 60 min of incubation with 50 nM of Topo I. When incubation was performed in the presence MgCl_2_, topoisomers with lower linking numbers were observed due to cleavage followed by religation. These results demonstrated that, as in other bacteria, the pneumococcal Topo I does not require Mg^2+^ for the cleavage step.

**Figure 8.**
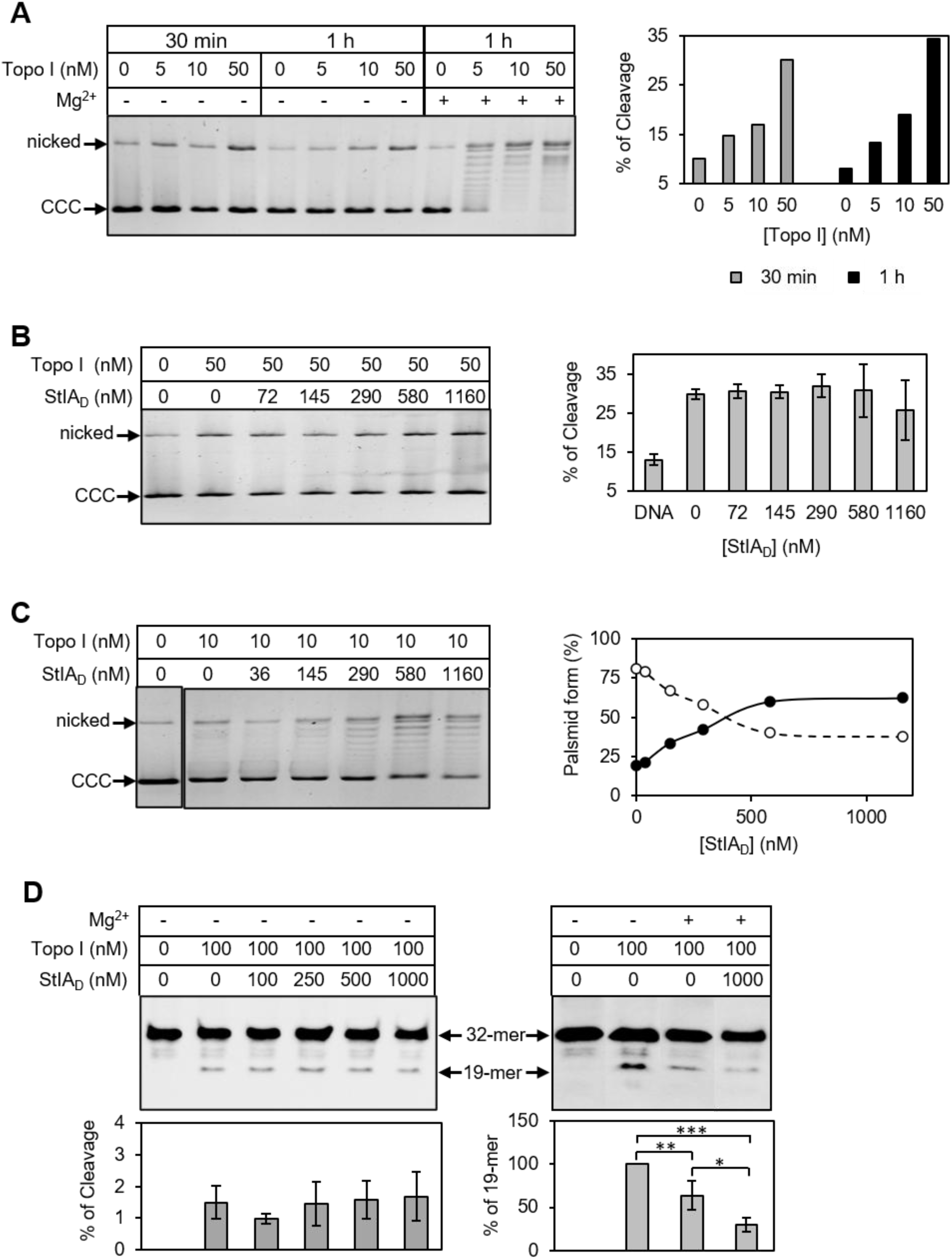
StIA specifically activates the intramolecular religation catalysed by Topo I. **(A)** Magnesium dependence of DNA cleavage or religation activities. CCC pBR322 (1 nM) was incubated with the indicated concentrations of Topo I in the absence or in the presence of 10 mM MgCl_2_. Reactions were analysed by agarose electrophoresis (left panel) and percentage of cleavage was quantified (right panel) as the amount of nicked plasmid relative to the total amount of plasmid in the well. **(B)** Effect of StIA_D_ on the cleavage activity of Topo I. 1 nM of pBR322 was incubated with Topo I at 4°C in the cleavage buffer, after 15 min, the indicated concentrations of StIA_D_ were added. Reactions were incubated for 1h at 37°C. A typical experiment is shown on the left and the percentage of cleavage (mean ± SEM, n=3) is represented on the right. **(C)** Effect of StIA_D_ on cleavage plus religation. First a cleavage reaction was performed as in (B). After 30 min of incubation at 37°C, 10 mM of MgCl_2_ was added to the reaction and incubation continued for other 30 min. The percentage of CCC (white circles) and topoisomers (black circles) were quantified (right panel). Reactions in A, B and C were analysed in 1.2% agarose gels, stained with ethidium bromide and quantified using Image Lab software. **(D)** Oligo-based cleavage (left) and religation (right) assays. TopoI–DNA cleavage complexes were generated in the absence of Mg^2+^ at 37 °C, followed by the addition of StIA_D_ to respective reactions, and subsequently Mg^2+^. Percentage of cleavage was quantified as the amount of 19-mer product relative to total DNA in the well. Religation was estimated by measuring the amount of 19-mer product relative to that in the absence of Mg^2+^. Three independent replicates were performed and the mean ± SD is plotted under each well. Statistical significance was analysed by *t*-Student test (*, *p* -value ≤ 0.1; **, *p* -value ≤ 0.05, *p* -value ≤ 0.01).

This property was then used to assess whether the StIA_D_ activates Topo I at the nicking step. To facilitate binding of Topo I to the DNA in a cleavage competent complex, we incubated pBR322 with Topo I at 4°C in the cleavage buffer prior to the addition of increasing concentrations of StIA_D_. Topo I exhibited a 30% of cleavage activity independently of the amount of StIA_D_ (Figure 8B). However, when 10 mM MgCl_2_ was added to the reaction to allow religation, DNA relaxation increases with StIA_D_ concentration (Figure 8C). This activation appeared to be independent of Topo I binding to DNA, as Topo I was preincubated with the DNA substrate before the addition of StIA. Therefore, StIA does not affect the DNA cleavage reaction and activation of Topo I is likely due to stimulation of the subsequent steps in the catalytic cycle.

The effect of StIA on the religation step was then evaluated using a 32-mer biotinylated oligonucleotide containing a strong cleavage site (STS) for *Mycobacterium tuberculosis* Topo I as substrate (Godbole et al., 2015). A standard cleavage assay performed in the absence of Mg^2+^ revealed that the pneumococcal Topo I was able to cleave the STS-containing oligonucleotide generating a 19-mer product. As expected, the presence of StIA_D_ did not affect DNA cleavage (Fig. 8D). For the religation assay, 1 µM of StIA_D_ was added to the reaction after 1h of cleavage, and religation was initiated by the subsequent addition of Mg^2+^. Intramolecular religation was monitored by quantifying the reduction in the 19-mer cleavage product relative to the condition without Mg^2+^ (Fig. 8D). In the absence of StIA_D_, addition of Mg^2+^ reduced the amount of the 19-mer product to 63.6 %, consistent with Topo I-mediated intramolecular religation. In the presence of StIA_D_, the level of the 19-mer product decreased further to 30.2 %, showing a 2.1-fold increase in religation efficiency. These results demonstrate that StIA_D_-mediated activation of Topo I occurs at the religation step of the catalytic cycle.

### 3.8. Protein modelling reveals distinct dimerization and Topo I interaction surfaces in StIA

The molecular interactions of StIA were analysed by structural modelling and protein-protein docking. A homodimeric structural model of StIA (M0) was obtained by ClusPro. The M0 model adopted a defined dimeric architecture with a concentrated intermolecular interface. The two StIA protomers buried a total surface area of 1,822.32 Å2, corresponding to approximately 911.16 Å2 per protomer. A total of 170 atom–atom contacts within 4 Å were identified, with a minimum intermolecular distance of 2.48 Å. The interface comprised 21 distinct residues, of which 17 were shared between the two protomer interfaces, indicating a spatially defined dimerization surface (Figure 9A, B and E)

**Figure 9.**
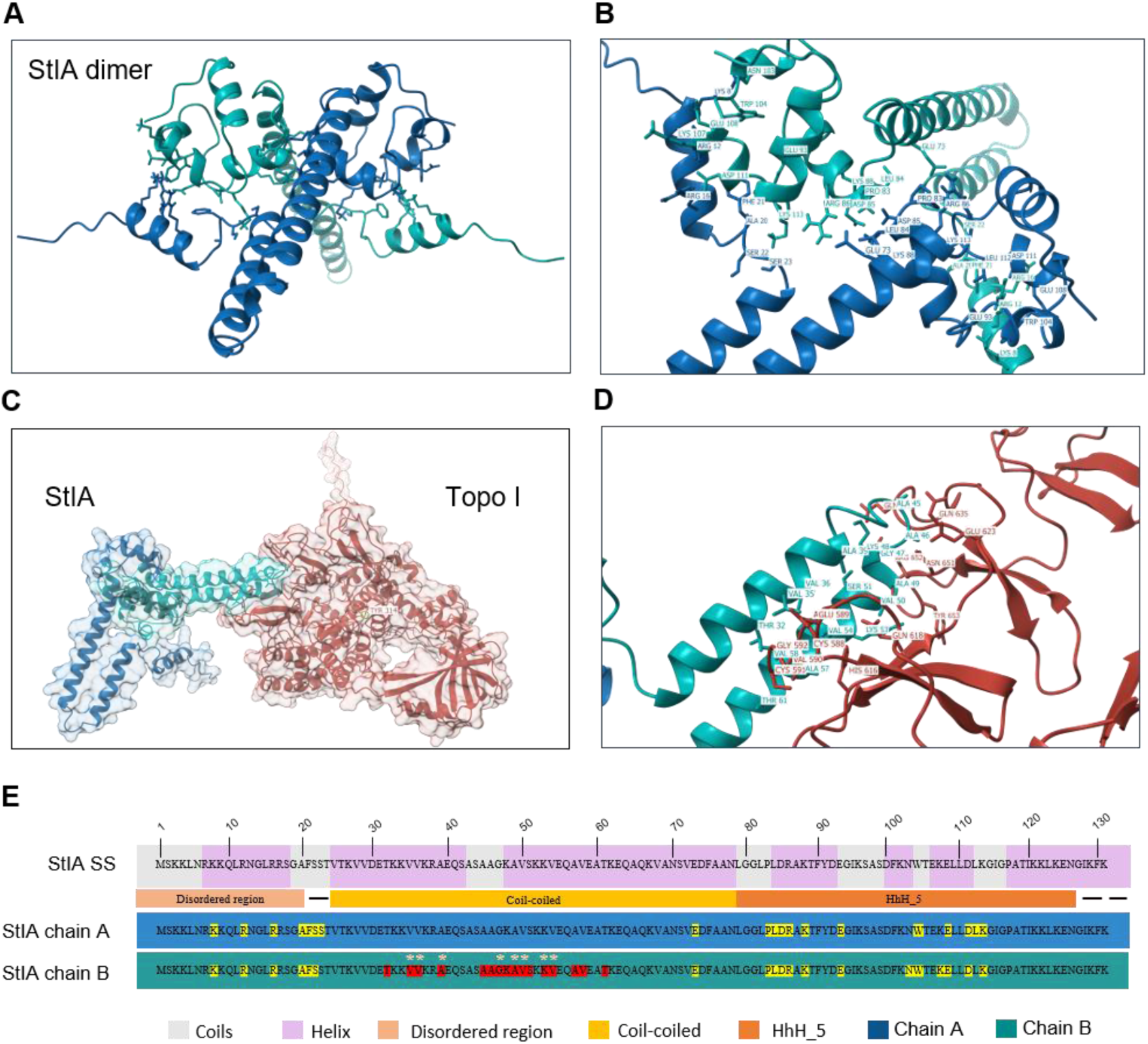
Exploratory structural analysis of StIA. **(A)** Highest-ranked model of the StIA M0 dimer generated using the ClusPro Multimer server. **(B)** Close-up view of the StIA M0 dimerization interface. Residues involved in intermolecular interactions between the two chains are highlighted shown. **(C)** Rank 1 prediction model of StIA – Topo I interaction obtained by PyDock. The catalytic Tyr314 of Topo I is indicated. **(D)** Contact residues at the StIA – Topo I interface. Intermolecular contacts were defined using a 4.0 Å heavy-atom distance cutoff. Contacting residues were identified using the open-source version of PyMol. **(E)** Structural and intermolecular interface map of StIA. The secondary structure of StIA is shown, with coils represented in gray and α-helices in pink. The predicted intrinsically disordered region, the coil-coiled and the HhH_5 domain are indicated below. The two StIA chains are displayed, highlighting residues involved in intermolecular interactions. Residues participating in dimerization are shown in yellow, whereas residues involved in Topo I interaction are shown in red. Asterisks indicate residues consistently predicted in different models to contribute to the StIA – Topo I interaction interface.

For StIA-Topo I interaction two independent docking models, pyDock and ClusPro, were used. The two approaches identified a recurrent central region of StIA as the principal candidate for interaction with Topo I. The three best ranked pyDock models yielded buried surface area values of 1,258.3, 1,187.7 and 855.6 Å2, respectively. pyDock Rank 1 and Rank 3 converged on a central StIA region encompassing residues 35–57, whereas Rank 2 represented an alternative mode centred more distally around residues 61–78. The pyDock Rank 1 model was selected as the representative pose for visualization of the StIA₂–Topo I interaction based on its favourable docking score and consistency with the alternative docking solutions (Figure 9C and D). This model was further supported by convergence with that obtained with ClusPro, which reproduced the same central StIA surface in its first two models.

Across the five principal models docking generated by pyDock and ClusPro, residues 35, 36, 39, 47, 49, 50, 53 and 54 occurred in four of five models. These residues define the strongest residue-level consensus and delineate a central StIA interaction surface centred approximately on residues 35–54, with additional recurrent contacts extending toward residues 57–58. Agreement between independent docking approaches is particularly informative because the two platforms use different sampling and scoring strategies. ClusPro Rank 3 produced a larger apparent buried surface area (2,900.5 Å2) and represents an alternative mode of interaction engaging both StIA monomers with a substantially different interaction geometry.

Importantly, the catalytic Tyr314 of Topo I showed a solvent-accessible surface area of 9.46 Å2 in Topo I and the same value after addition of StIA.

## 4. Discussion

In *S. pneumoniae*, Topo I is the primary DNA relaxation enzyme, and alterations in its activity or cellular abundance can significantly perturb chromosome topology (Ferrándiz et al., 2016; García-López et al., 2023b). In this study, we identify StIA as a previously uncharacterized protein that specifically enhances the activity of pneumococcal Topo I. The *stIA* gene is highly conserved among all sequenced pneumococcal genomes and is also present in other *Streptococcus* species, where it remains annotated as hypothetical protein of unknown function. Alterations in StIA levels did not affect bacterial growth in the presence of the gyrase inhibitor NOV. In contrast, deletion of *stIA* impaired growth under subinhibitory concentrations of the Topo I inhibitor SCN, whereas StIA overexpression increased resistance to this drug (Figure 3). Consistent with these phenotypic observations, our biochemical analyses demonstrate that StlA specifically stimulates Topo I activity without affecting gyrase (Figure 4).

Stimulation of Topo I by StIA may occur through at least two non-mutually exclusive mechanisms. First, StIA may directly interact with Topo I to enhance its catalytic activity. Alternatively, it may bind DNA and alter its conformation, thereby generating a more favourable substrate for the enzyme. Both mechanisms have precedents in bacterial systems. In *M. tuberculosis*, a DNA glycosylase and a ribokinase have been reported to modulate Topo I through direct protein-protein interactions (Huang and He, 2010; Yang et al., 2011, 2012). By contrast, the *E. coli* single-stranded DNA binding protein (SSB) stimulates Topo I without detectable physical interaction by promoting its binding to DNA (Sikder et al., 2001). A more complex mode of regulation is exerted by HU, the major bacterial NAP, which stimulates Topo I activity at low concentrations through direct interaction with the enzyme, whereas at higher concentrations it inhibits Topo I and other topoisomerases by occluding DNA (Ghosh et al., 2014). Furthermore, the ability of HU to constrain DNA topology has also been proposed to indirectly modulate Topo I activity by altering the architecture of its DNA substrate (Bensaid et al., 1996).

Our data show that StIA binds dsDNA in a sequence-independent manner, consistent with the presence of the predicted C-terminal HhH domain. When incubated with long dsDNA molecules *in vitro*, StIA forms DNA-protein complexes that fail to enter the gel during electrophoresis, suggesting the assembly of higher-order oligomeric structures. Similar higher-order oligomers were detected *in vivo*, where they co-eluted with the nucleoid fraction. Thus, although free StIA predominantly exists as a dimer, DNA binding appears to promote higher-order assemblies that coat the DNA. This kind of binding could alter DNA conformation and/or reduce its effective charge, thereby influencing Topo I activity.

However, the H_6_-StIA fusion protein exhibited a markedly reduced ability to stimulate Topo I (Figure 4) despite displaying DNA-binding activity comparable to that of StIA_D_ (Figure 6), suggesting that Topo I activation is largely independent of DNA binding. Furthermore, StIA_D_ bound CCC and RC plasmids, the preferred substrates of Topo I and gyrase, respectively, with similar efficiencies, whereas its stimulatory effect was restricted to Topo I. Together, these observations argue against a mechanism in which StIA activates Topo I primarily by altering substrate conformation. Nevertheless, subtle structural differences between H_6_-StIA and StIA_D_ were detected by size exclusion chromatography, and additional effects on DNA architecture cannot be discarded. Indeed, although both proteins bound to short oligonucleotides with similar efficiencies, differences became apparent when larger dsDNAs were used. H_6_-StIA and StIA_D_ bound to dsDNA with comparable stoichiometry, but StIA_D_ exhibited a greater capacity to assemble higher-order oligomeric complexes, a property that could ultimately contribute to Topo I activation.

Alternatively, StIA may stimulate catalytic activity of Topo I through direct physical interaction by recruiting the enzyme to the DNA, stabilizing the Topo I-DNA complex, or facilitating one or more catalytic steps of the reaction. Our biochemical studies demonstrate that StIA directly interact with Topo I *in vitro* forming a stable StIA – Topo I complex. Furthermore, we have shown that StIA does not affect the DNA cleavage activity of Topo I but enhances subsequent steps of the relaxation reaction and this stimulatory effect is independent of Topo I DNA-binding. Rather, StIA specifically promotes the intramolecular religation catalysed by Topo I. A related mechanism has been described for *M. tuberculosis* HU, which specifically activates Topo I by enhancing the strand-passage activity without affecting either DNA cleavage or religation steps (Ghosh et al., 2014). Collectively, these findings strongly support a model in which StIA activate Topo I through a direct protein-protein interaction, thereby enhancing the religation step of the catalytic cycle.

Structural analysis supports a model in which StIA forms a defined homodimer and provides a coherent framework for interpreting the experimentally observed interaction between StIA and Topo I. Independent pyDock and ClusPro calculations converged on a central StIA surface, with eight consensus interacting residues. This interacting surface is spatially separated from the catalytic Tyr314 of Topo I, which shows the same solvent-accessibility before and after StIA association. This structural model is particularly relevant to the biochemical observation that StIA stimulates Topo I. Topo I undergoes coordinated DNA binding, single-strand cleavage, strand passage and religation, and their functional cycle depends on conformational rearrangements (Garnier et al., 2018; Dasgupta et al., 2020). StIA engaging a non-catalytic surface of Topo I could therefore influence Topo I activity without directly occupying the catalytic centre. Moreover, structural predictions identified N-terminal and intermediate surfaces involved in dimerization and Topo I interaction, respectively. This is consistent with the differences observed on dsDNA binding and Topo I stimulation upon the H₆-tag removal and further support the relevance of the N-terminal region of StIA to Topo I modulation. Future targeted mutagenesis of recurrent StIA residues, particularly within the 35–54 region, combined with interaction and enzymatic assays, may help to assess the functional relevance of the predicted interface.

To date, only two proteins have been shown to modulate Topo I activity in *S. pneumoniae*: HU, which has been proposed to constrain the DNA relaxed by Topo I (Ferrándiz et al., 2018), and StaR, which stimulates Topo I activity (de Vasconcelos Junior et al., 2023). The former appears to act *via* DNA binding, whereas the latter modulates Topo I through direct protein-protein interaction. The amount of both proteins correlates with the levels of Sc and their expression is downregulated under inhibitory concentrations of NOV, when chromosomal DNA becomes relaxed. By contrast, StIA expression is not regulated by Sc, and its cellular levels remain unchanged following NOV treatment. Similar to StaR, StIA directly interacts with and specifically activates Topo I, while, like HU, it also binds dsDNA and promotes the formation of high-molecular-weight nucleoprotein complexes. We demonstrate that StIA enhances the post-cleavage religation step of the catalytic cycle but given its strong capacity to bind dsDNA, additional stimulatory effects due to substrate modification cannot be discarded. Collectively, these findings support the existence of three proteins that modulate the pneumococcal Topo I activity through distinct mechanisms, as reported in other bacteria. Topoisomerases are essential enzymes in the maintenance of DNA topological homeostasis, and their functional cooperation with accessory proteins, although not essential for viability, is likely to play an important role in preserving chromosome topology. The activity of gyrase and Topo IV are also regulated by accessory proteins in several bacterial species, but comparable mechanisms have not yet been identified in *S. pneumoniae*.

Although no Topo I-targeting antibiotics are currently used in clinical practice, the essential role of this enzyme makes it an attractive target for the development of new antibacterial agents. Our results show that StIA modulate susceptibility to SCN *in vivo*. SCN, a compound discovered by our group, specifically inhibits the DNA cleavage reaction catalysed by bacterial Topo I and suppress the growth of *S. pneumoniae* (García et al., 2011; García-López et al., 2023a) and *M. tuberculosis* (García et al., 2018). Moreover, SCN exhibits greater bactericidal activity than FQs, against both planktonic bacteria and biofilms (Valenzuela et al., 2020) and is effective against FQ-resistant pneumococcal isolates in a murine sepsis model (Tirado-Vélez et al., 2021). Therefore, SCN, and other Topo I inhibitors represent promising alternative treatments for pneumococcal infections, including those caused by FQ-resistant strains. More broadly, a detailed understanding of the factors that regulate Topo I activity may provide foundation for the development of novel therapeutic strategies targeting this essential enzyme.

Moreover, StIA appears to be associated with the pneumococcal nucleoid, a membrane-less intracellular compartment that is maintained in a state of phase-separation from cell cytoplasm through the coacervation of NAPs with DNA and/or RNA. It has been proposed that NAPs involved in the maintenance of such state should be highly conserved in bacteria, highly abundant, capable of binding nucleic acids in a sequence-independent manner and contain intrinsically disordered regions that would promote phase separation. In *E. coli*, two NAPs, HU and Dps, fulfil these criteria. Both proteins undergo complex coacervation with DNA and RNA *in vitro*, resulting in the formation of liquid-liquid phase-separated condensates, a behaviour proposed to be a model for the nucleoid organization *in vivo* (Gupta et al., 2023). In addition, both proteins exhibit a high degree of multimerization, a property that correlate with their ability to form condensates with DNA. Similarly, StIA is a highly conserved, sequence-independent DNA-binding protein that contains a predicted intrinsically disordered region and assembles into high-order oligomers upon DNA binding. These characteristics suggest that StIA may contribute to establishment and/or maintenance of the phase-separation state of the pneumococcal nucleoid.

## Author contribution

AVJ performed the experimental work, the structural modelling and participated in data interpretation. PH carried out the *in vitro* cross-linking assays. MA participated in the experimental work. AGC got funding and administered the project. AGC and MA participated in data interpretation, conceived, designed, supervised the study and wrote de manuscript. All authors participated in the correction of the manuscript. The manuscript has been approved by all authors for publication.

## Acknowledgements

We thank María José Ferrándiz (CNM, ISCIII, Spain) for providing antibodies against HU. During the preparation of this work, the authors used ChatGPT (OpenAI, free version) and Microsoft Copilot to improve the grammar, language, and readability of the manuscript text. After using these tools, the authors reviewed and edited the content as needed and took full responsibility for the content of the published article.

